# Complex Assembly and Activity States as Multifaceted Protein Attributes Explaining Phenotypic Variability

**DOI:** 10.1101/2025.09.25.678633

**Authors:** George Rosenberger, Peng Xue, Isabell Bludau, Claudia Martelli, Evan Williams, Ben C. Collins, Andrea Califano, Yansheng Liu, Ruedi Aebersold

**Author notes:** Equal Contribution (George Rosenberger, Peng Xue).

## Abstract

Cell function studies primarily focus on measuring overall molecular abundances while often overlooking critical clues—including protein modifications and molecular interaction networks—that critically determine the functional properties of the cell. In prior work, we introduced a suite of methods to reveal context-specific transcription factor-gene regulatory networks, kinase-substrate networks, and protein interaction networks and leveraged them to gain deeper insights into transcriptional regulation and signal transduction. However, the complex interdependencies between these networks are still elusive. To address this challenge, we introduce a multi-omics framework, aimed at harnessing measured or inferred protein activity in context-specific networks, which yields deeper functional insights into mechanisms underlying molecular phenotypes, compared to protein abundance alone. As proof of concept, we utilized progressively differentiated instances of HeLa CCL2 and Kyoto cell lines to explore the role of protein complexes and interactions in cell doubling time and susceptibility to *Salmonella* Typhimurium infection. Notably, this analysis underscores the pivotal role of protein interaction networks in linking molecular profiles to phenotypic outcomes, thus providing a highly generalizable framework for multi-omics dataset analysis.

## Introduction

Phenotypes arise from the complex biochemical states of cells and tissues. These states, in turn, reflect the aggregate and interplay of genes, transcripts, proteins, lipids, and metabolites, a network intricately woven by the central dogma of molecular biology [1]. Understanding how phenotypes connect to these underlying molecular landscapes and the molecular mechanisms they constitute remains a fundamental pursuit in both basic and biomedical research. The advent of systems biology [2] and the rapid evolution of molecular profiling techniques, especially multi-omics technologies, offer an unprecedented opportunity for the holistic exploration of biological systems. However, as multi-omics profiling technologies become more accessible, interpretation of their associated molecular profiles remains challenging. Many data integration methods struggle with the need to identify causal rather than associational links between molecular abundances and cell function [3]. In particular, the role of proteins, essential in controlling biological processes, exemplifies this complexity, as their function and activity depend on intricate mechanisms of molecular state-dependent interactions.

To address these challenges, we and others have developed network-based approaches that integrate prior and *de novo* knowledge, including generalized or context-specific molecular interaction networks generated by computational analysis of large-scale data repositories or using experimental techniques [3]. Validated uses of such approaches include, for instance, the assessment of transcription factor (TF) and co-factor (co-TF) activity from gene regulatory networks—as exemplified for instance by the Virtual Inference of Protein activity using Enriched Regulon analysis, VIPER [4]—or assessment of kinase/phosphatase activity using signal transduction networks and phosphoproteomic profiles—as exemplified by algorithms such as Kinase Substrate Enrichment Analysis (KSEA [5]), PHOsphorylation NEtworks for Mass Spectrometry (PHONEMeS [6]), or Virtual Enrichment-based Signaling Protein-activity Analysis (VESPA [7]). To further extend the ability of these methodologies to build quantitative, mechanistic models of cell behavior [8], we recently developed a high-resolution protein co-elution profiling approach (SEC-SWATH-MS [9, 10]) and the Size-Exclusion Chromatography Algorithmic Toolkit (SECAT [11]). These tools assess context-specific protein network states using metrics that represent protein complex abundance and stoichiometry based on quantitatively characterized protein-protein interactions (PPIs). SECAT’s focus on protein network state, rather than static protein complexes, allows capturing dynamic state/condition-dependent changes in protein complex formation and stoichiometry, making it ideally suited to characterizing protein complex dynamics from multi-omics profiles.

In this study, we explore multiomic, network-based inference of protein activity as a more effective approach to unravel phenotype-associated molecular mechanisms, compared to direct transcriptomic or proteomic abundance analysis. For this purpose, we leveraged previously generated data on the molecular and phenotypic heterogeneity of HeLa cells, using 14 HeLa samples gathered from 13 laboratories [12]. Cells were subjected to multi-omics profiling to assess quantitative effects of genomic variability on the transcriptome and proteome, including protein turnover. A key discovery was the robust correlation between “proteotypes”—i.e., the contextual state of the proteome—and cell phenotypes, particularly cell doubling time and susceptibility to *Salmonella* Typhimurium infection in the HeLa panel. The study led to three hypotheses that were tested in the current work: First, protein complex stoichiometry represents an effective buffering mechanism against copy number (CNV) and transcriptional heterogeneity. Second, protein function is more closely related to molecular mechanisms and cellular phenotype than protein abundance, and third, that multi-omics data provide complementary information to connect molecular profiles to phenotypes.

Our study demonstrates the importance of characterizing context-specific, dynamic protein complex composition as part of the proteotype and the potential value of the proposed methodology for future multi-omics-based studies.

## Results

### Outline and graphical abstract

To investigate the buffering effects of protein complex stoichiometry against CNV and transcriptional variation, we first measured the two most distant instances of HeLa Kyoto and CCL2 cell lines by SEC-SWATH-MS to obtain high-resolution protein complex profiles (Fig. 1a). Quantitative assessment of the effects of copy number variation (CNV) and mRNA abundance on monomeric and assembled protein complex subunits allowed us to draw generalized conclusions about protein subunit buffering in cellular systems.

**Figure 1.**
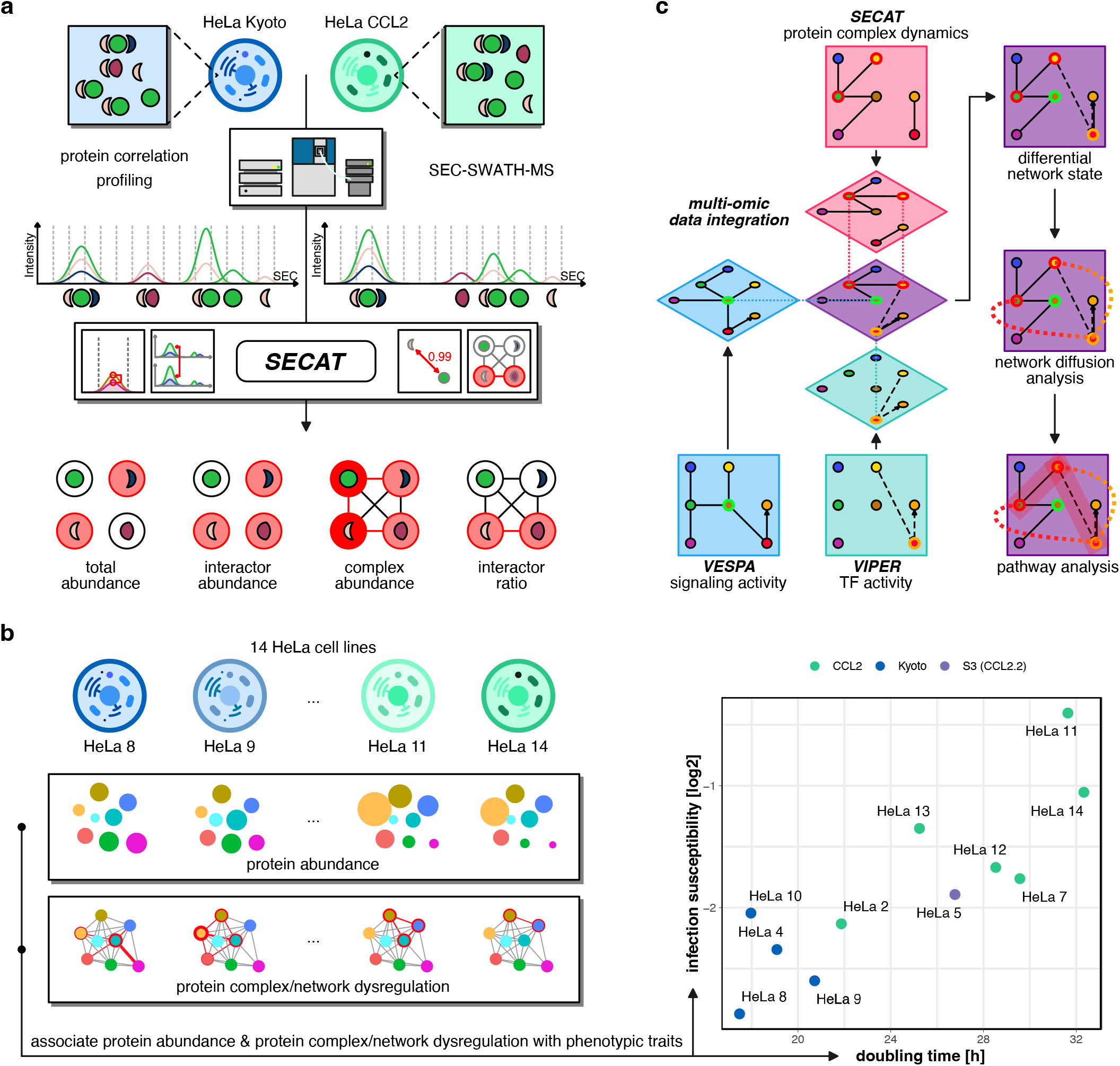
Methodological overview. **a)** Protein complex profiles were generated by SEC-SWATH-MS and then analyzed using the Size-Exclusion Chromatography Algorithmic Toolkit (SECAT). SECAT uses a STRING protein-protein interaction network (PPIN) as reference to generate a context-specific network and to infer protein attributes for complex abundance and stoichiometry. **b)** Association of protein abundance and protein complex/network dysregulation with two phenotypic traits of HeLa cell lines, doubling time and infection susceptibility. **c)** Differential transcription factor (TF) activity was inferred using VIPER, differential signaling activity was inferred using VESPA, and protein complex attributes were obtained from the SECAT analysis. The combined results provide an overview on the connectivity of proteins and their differences between the two HeLa subtypes. Network diffusion analysis makes use of the context-specific PPIN and GRN to assess the effects of upstream alterations on downstream layers.

We then used the 14 HeLa cell line panel to match proteotypes to two quantitatively and experimentally assessed phenotypic traits (Fig. 1b): Cell doubling time and susceptibility to *Salmonella* Typhimurium infection [12]. For this purpose, we contextualized protein-protein interaction networks and used them to infer protein complex dynamics. Our analysis demonstrates that protein activity is more closely related to phenotypic traits than protein abundance, although the underlying data is identical.

Finally, we conducted a differential comparison of HeLa CCL2 vs. Kyoto using the inferred protein activities of VESPA, VIPER, and SECAT. A network diffusion approach elucidated the functional dependencies between the two cell lines, ranging from altered signaling and transcriptional programming to reorganization of the protein complexes and macromolecular structures involved in the organization of the extracellular matrix, which are directly responsible for previously observed phenotypes (Fig. 1c).

### Quantitative effects of copy number variation and mRNA abundance on protein complex assembly

Approximately 60% of protein mass in cell extracts under mild conditions is observed in complexed form [10], with complex subunits typically synthesized at similar levels [13]. Since complex formation influences function, we hypothesized that CNV and mRNA variation affect assembled and monomeric protein fractions differently. Our previous study showed that proteins known to be part of complexes are less impacted by such variation, suggesting that monomeric subunits may buffer synthesis variability, particularly under abnormal gene dosages [12]. The complex-resolved data from this study now allowed us to explore these findings in greater detail.

We first tested whether complex stoichiometries represent an effective buffering mechanism contributing to the divergence of the effects of copy number variation (CNV) at the transcript and protein level. This question is of significant functional importance because it can be assumed that the functions of complex-bound and monomeric forms of a protein are functionally different. We leveraged our previously developed size exclusion chromatography (SEC)-based approach SEC-SWATH-MS [10] in combination with the SECAT algorithm [11] (Fig. 1a, Methods) to determine the context-specific PPI maps of two representative HeLa Kyoto and HeLa CCL2 cell lines and carried out protein complex profiling in three replicates for each cell line (Methods). In total, the quantitative matrix encompassed 7,175 proteins with 89,065 unique peptides assigned to these proteins (1% global context FDR; only proteotypic peptides), detected from a total of 420 LC-MS/MS runs (Methods). SECAT was then used to generate context-specific PPI networks, to infer protein-level attributes inducing changes in protein complex composition and abundance, and to quantify the distribution of complex-bound and monomeric fractions of detected proteins between the two conditions (Methods, Supplemental Fig. 1-2, Supplemental Data 1).

First, we deconvoluted protein abundance into monomeric and assembled states by SEC-SWATH-MS. Then, we combined these results with available, cell line-matched CNV, mRNA and protein abundance data from our prior study to investigate the effect of assembly state on protein abundance buffering (Fig. 2, Supplemental Data 2). We found that within-gene differential CNV, mRNA abundance and total protein abundance levels are correlated, as previously shown [12] (CNV-protein: Spearman’s ρ: 0.248, original study: 0.217; mRNA-protein: Spearman’s ρ: 0.514, original study: 0.510). The quantitative SECAT metrics, however, provide strong support for the monomer “buffering” hypothesis (Fig. 2). Here, proteins present exclusively in either monomeric or assembled conformation are called “single state”, whereas proteins present in both monomeric and assembled conformation are called “paired state” (Fig. 2a).

**Figure 2.**
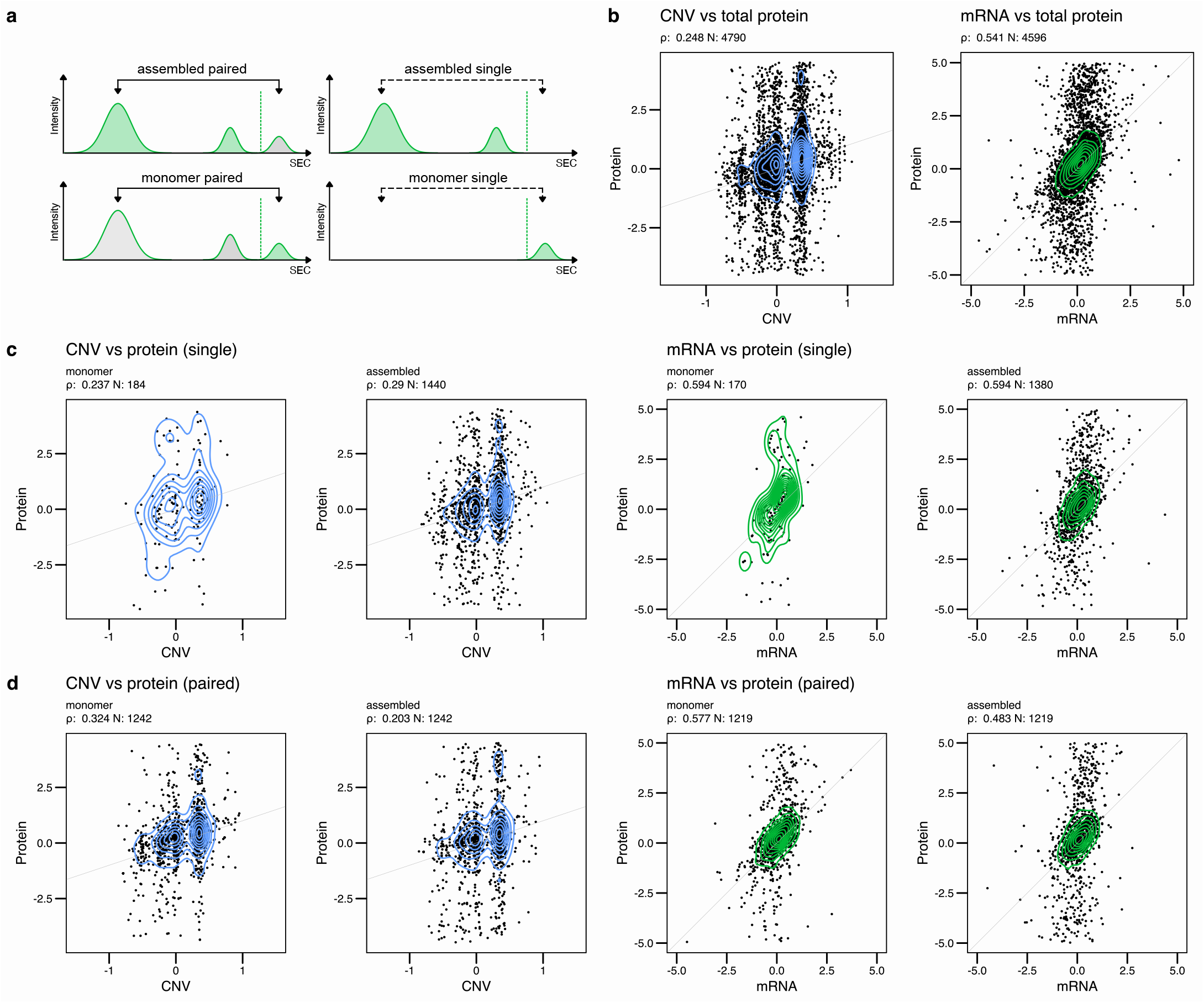
Dependency of monomer and assembled levels on copy number variation (CNV) and mRNA abundance. **a)** Definition of paired and single monomer and assembled protein fractions. **b)** CNV and mRNA abundance vs. total protein abundance. **c)** CNV and mRNA abundance vs. single monomeric and assembled protein abundance. **d)** CNV and mRNA abundance vs. paired monomeric and assembled protein abundance.

“Single state” monomeric and assembled fractions are similarly correlated to CNV and mRNA abundance (monomer/assembled: Spearman’s ρ: 0.237/0.29 (CNV-protein); 0.594/0.594 (mRNA-protein); CNV-protein p-value: 0.236; mRNA-protein p-value: 0.499, Methods). In contrast, for the “paired state” proteins, monomeric fractions emerged as significantly more correlated to CNV and mRNA abundance than the assembled fractions (monomer/assembled: Spearman’s ρ: 0.324/0.203 (CNV-protein); 0.577/0.483 (mRNA-protein); CNV-protein p-value: 0.00059; mRNA-protein p-value: 0.0006, Methods). This observation is consistent with the assumption that heteromeric complex subunits are individually regulated, at least in some cases. Specifically, if the cognate binding partners of a subunit are not available—due to CNV/mRNA variability—this would result in “spare” monomers, thus reducing the assembled fraction’s overall correlation. Thus, integrated analysis of protein complex, CNV, and mRNA profiles suggests that proteins in assembled conformations are buffered from CNV and mRNA abundance variation compared to their monomers.

Although mRNA and protein abundance is weakly albeit significantly correlated at steady state [12, 14, 15], decomposing the factors contributing to this effect across different proteins is challenging. Here, we investigate the issue using protein complex-resolved profiles, thus providing direct evidence of stoichiometry-related buffering. In addition to gene dosage and transcription rates, other contributing factors may include splicing or proteasomal-mediated protein turnover [16]. Further, protein activity, as opposed to abundance, is also highly dependent on additional factors such as post-translational modification state, subcellular localization and protein complex interaction partners. Thus decomposing protein abundance into different activity states might be more informative.

### Association of protein abundance and protein complex/network dysregulation with phenotypic variability

A key challenge in proteomic biomarker studies is translating statistically significant protein abundance signatures, which effectively distinguish cellular or disease states, into clinically actionable insights. The difficulty lies in linking these proteins to their fundamental molecular mechanisms. Drawing inspiration from transcriptomics, where network-based cell state analyses consistently yield more robust and functionally relevant biomarkers than direct transcript profiles [4], we hypothesize a parallel advantage. Specifically, we suggest that protein interaction states—the condition-specific patterns of protein-protein interactions—could offer more mechanistic insights compared to isolated protein abundances.

To address this question, we assessed whether phenotypic trait variability may be better explained by differences in protein abundance or protein interaction states, and whether the latter may be more effectively linked to molecular mechanism. For this purpose, we focus on two established quantitative traits that have shown substantial variability in HeLa samples, their doubling time, and their susceptibility to *Salmonella* Typhimurium infection [12].

To generate context-specific protein-protein interaction (PPI) networks and assess protein complex abundance directly from whole, unfractionated proteome profiles, we developed STELLA (Signature Transformation Enhanced Local Likelihood Analysis). STELLA assembles HeLa sample-specific PPI networks by integrating the CORUM [17] database (focused on protein complexes) and the STRING [18] database (encompassing both complex and transient interactions) with VESPA’s network pruning module [7], leveraging individual protein abundance profiles (Methods).

We then utilized the DeMAND [19] algorithm, a network-based approach identifying proteins with dysregulated interactions, to quantify protein complex (1,414 proteins) and protein network (4,966 proteins) dysregulation across experimental conditions for each cell line (Fig. 1b, Methods, Supplemental Data 3). These metrics, which assess changes in protein interactions within either complexes or broader networks, are anticipated to provide more potent predictions of protein function than simple abundance profiles.

Following established biomarker discovery frameworks, we generated phenotype-specific signatures for each tested data modality (mRNA transcript, protein, and phosphopeptide abundance) and for their corresponding inferred features, including transcription factor activity, kinase/phosphatase activity, and complex/network dysregulation. We performed recursive feature elimination using bootstrapped random forest regression, optimizing for Root Mean Square Error (RMSE). This approach provided variable importance estimates for the identification of molecular determinants linked to the phenotypes (Methods). The results showed comparable predictive performance across all data types, achieving R^2^ values between 0.4 and 0.6 using the top 128 features, consistent with the RMSE profiles (Fig. 3b, Supplemental Data 4).

**Figure 3.**
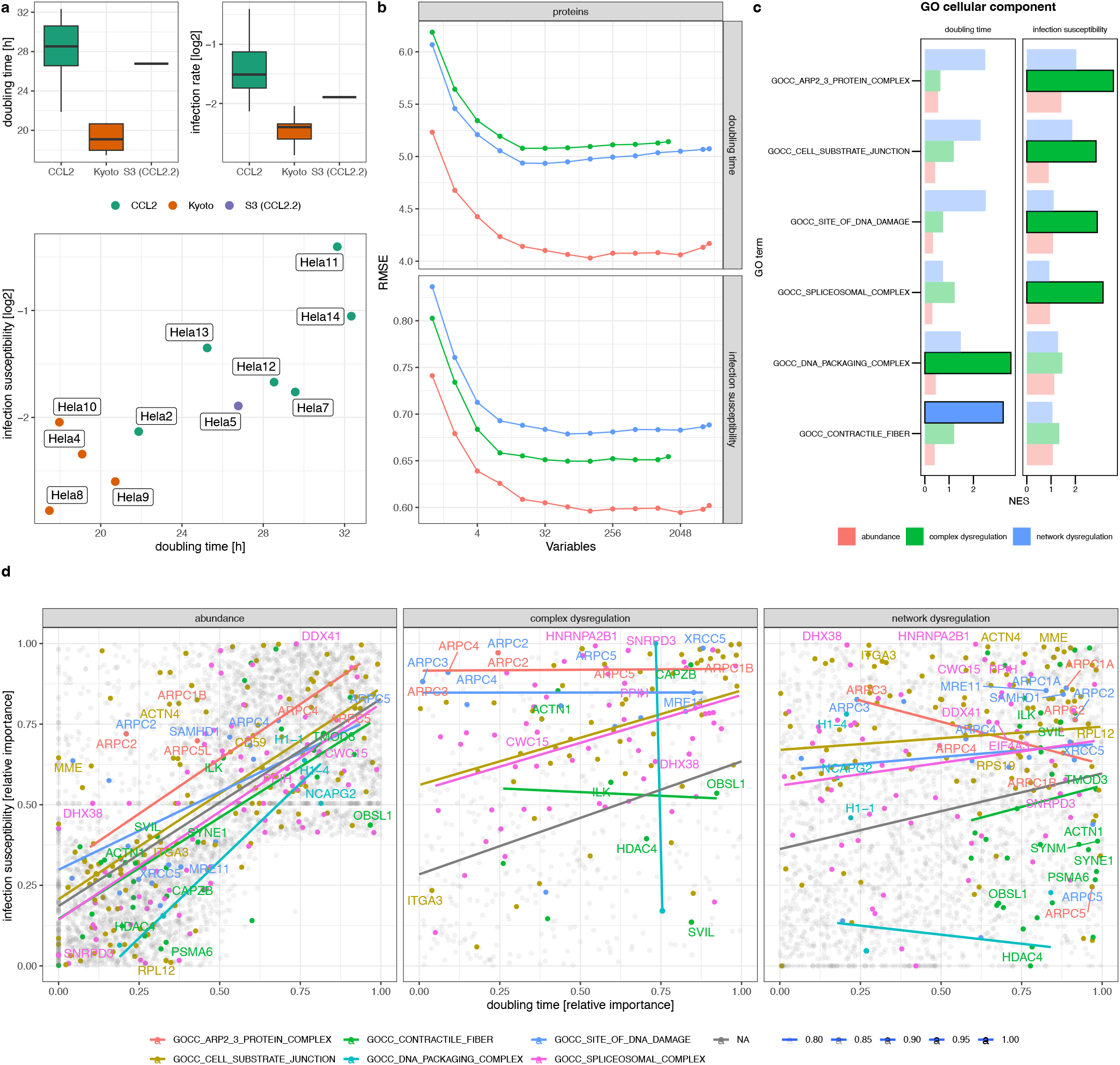
Association of protein abundance and protein complex/network dysregulation with phenotypic variability. **a)** Quantitative phenotypes characterizing cell doubling time and *Salmonella* Typhimurium infection susceptibility [12]. **b)** Recursive feature selection identified the best features according to RMSE. **c)** GO cellular components indicate different association of selected protein abundance and protein complex/network dysregulation features. Opaque categories are significant (BH-adj. p-value < 0.05). **d)** Correlation plots display relative feature importance between doubling time vs. infection susceptibility phenotypic traits. Strong correlation for abundance indicates lower association with a particular trait, whereas complex and network dysregulation associate features of related gene sets specifically with traits.

Although all individual data modalities showed similar predictive performance for the two phenotypes, we hypothesized that network states would be better suited to identify proteins associated to phenotype presentation, compared to protein abundance signatures alone. To test this, we performed Gene Set Enrichment Analysis (GSEA) [20, 21] using a Gene Ontology (GO) cellular component database to evaluate which modalities select features more closely associated with underlying mechanisms. The results revealed notable differences between protein abundance and protein complex or interaction network dysregulation. For “infection susceptibility,” complex dysregulation features were significantly enriched for GO terms related to the ARP2/3 complex, spliceosomal complex, cell-substrate junctions, and DNA damage sites—enrichments that were not significant by protein abundance analysis (Fig. 3c). In contrast, predictive features for “cell doubling time” showed enrichment for DNA packaging complexes and contractile fibers, distinct from those observed for infection susceptibility.

Interestingly, when comparing the association of features selected by doubling time vs. infection susceptibility analysis, a distinct protein abundance and complex/network dysregulation pattern emerged (Fig. 3d). Whereas differential abundance of the same protein was frequently nominated for both traits (Spearman’s ρ=0.66), complex, and network analyses identified different feature dysregulation in the two phenotypes without significant correlation between the selected features (Spearman’s ρ=0.37, p=1.45e-21 and ρ=0.23, p=9.26e-115 by one-tailed Spearman test), as shown by proteins in statistically significant GO terms (Fig. 3d). For instance, the Arp2/3 protein complex subunits ARPC1B, ARPC2, ARPC3, ARPC4 and ARPC5 are nominated among the most dysregulated in complex formation for infection susceptibility but not for doubling time, yet they are far less significant in terms of protein abundance. Similarly, proteins associated with GO term “contractile fiber” are nominated by network dysregulation analysis for the doubling time phenotype, yet they are also not statistically significant based on abundance. Many of these proteins, such as ILK, SVIL, SYNE1, PSMA6, OBSL1, and HDAC4 are closely associated with cell cycle regulation, especially through their molecular interactions. HDAC4, for example, a protein known to inhibit the activity of CDK1 [22]—a key regulator of cell cycle—only emerged as significant by protein complex/network dysregulation analysis but not by protein abundance analysis.

These results are supported by our previous observation of morphological differences between HeLa cell lines, especially in terms of altered actin structures, which may also be associated with the variation in infection susceptibility [12]. Although the small number of samples limit the generalizability of the conclusions that can be drawn from this analysis, the results suggest that in cases where molecular function is primarily mediated by interactions, assessment of contextual interaction dysregulation can be more informative than abundances alone.

### Differential kinase/phosphatase, transcription factor activities and protein complex dynamics of HeLa Kyoto vs. CCL2 cell lines

We next investigated whether protein activity, as inferred from multi-omic profiles, offer complementary and synergistic insights, surpassing individual contributions, to differentiate functional module states between HeLa Kyoto and CCL2 cell lines.

To infer differential kinase/phosphatase activities between the cell lines tested, we used the multi-sample VESPA [7] (msVESPA) algorithm and a contextualized signaling network covering 549 kinases and phosphatases (Methods, Supplemental Data 5). Using this approach and the previously published phosphoproteomic profiles of HeLa Kyoto vs. CCL2 cell lines [23] (6 cell lines each, measured in triplicates), we identified 71 out of 525 kinases and phosphatases to be differentially active (BH-adj. p-value<0.05, Fig 4, Supplemental Fig. 3). Gene Set Enrichment Analysis (GSEA) using the msVESPA normalized enrichment score (NES) identified metabolic kinases and phosphatases as the only significantly enriched gene set (adj. p-value<0.05, Fig. 4a). However, the most differentially active enzymes included critical kinases involved in vesicular transport (PTPN23), extracellular matrix organization (EEF2K, LMTK2, TRPM7, ABL1, PPM1F, LIMK1), the mitogen-activated protein (MAP) kinase cascade (MAPK13, MAP2K4, PRKCZ, FLT1, TYRO3, DUSP1), phosphatidylinositol 3-kinases (PI3K) signaling (NTRK2, NTRK3, FER, PDGFRA), Wnt signaling (CSNK1E, ROR2), JAK-STAT (JAK2) and others (Fig. 4b).

**Figure 4.**
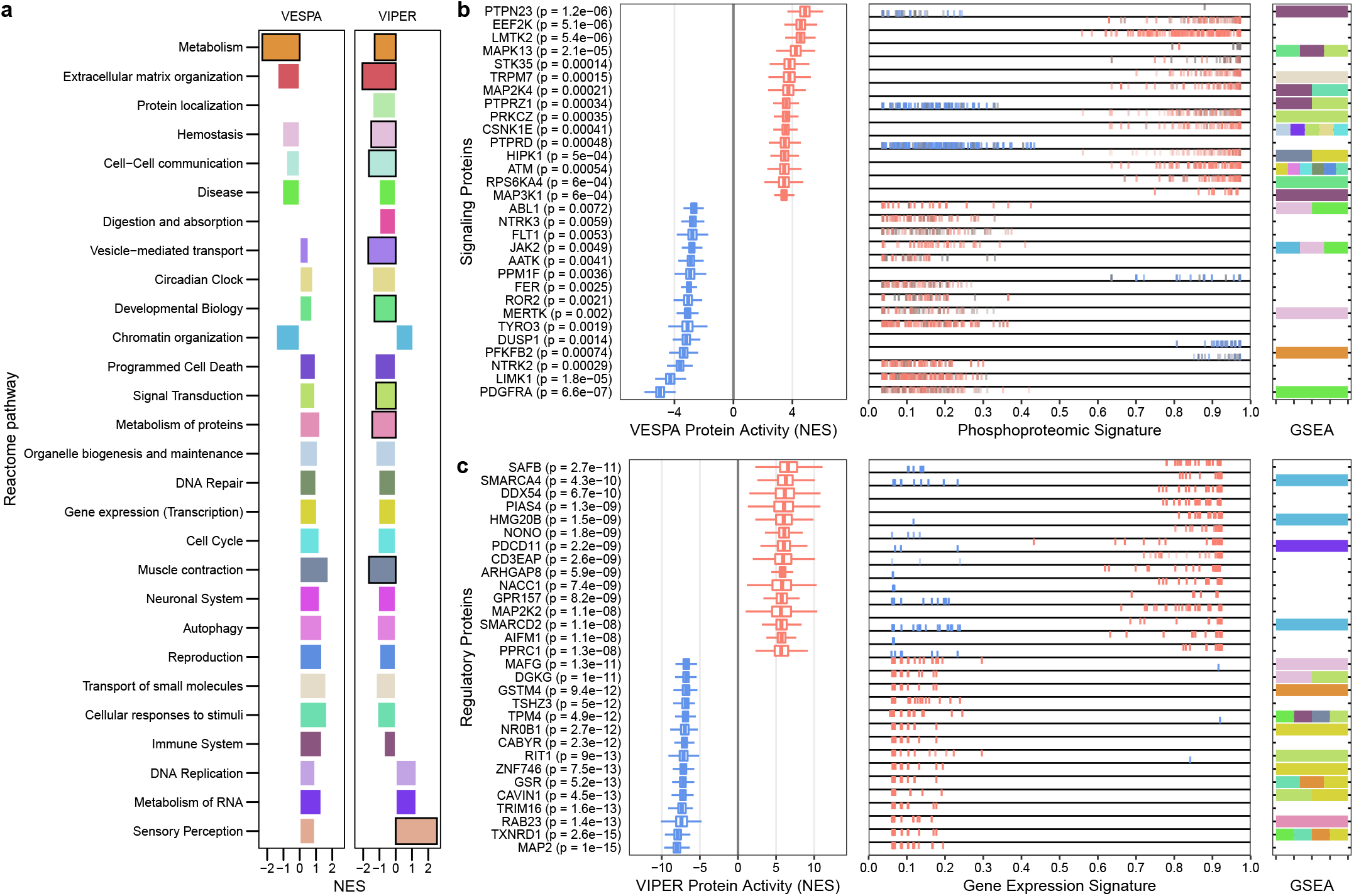
Differential transcription factor, kinase and phosphatase activities of HeLa Kyoto vs. CCL2 cell lines. **a)** Gene set enrichment analysis (GSEA) against the Reactome top pathways indicates differences between the cell line subtypes. Bold highlighted categories are significant (BH-adj. p-value < 0.05). **b)** The top 30 signaling proteins with differential signaling activity between the two conditions are depicted. **c)** The top 30 regulatory proteins with differential gene regulatory activity between the two conditions are depicted.

In parallel, to infer differential TF activities, we applied multi-sample VIPER [4] (msVIPER) analysis using a metaVIPER [24] contextualized gene regulatory network comprising 5,794 genes, generated from our previously published transcriptomic profiles for 6 HeLa Kyoto and 10 HeLa CCL2 cells (Methods, Supplemental Data 6).

Specifically, msVIPER was used to infer the activity of 5,950 TF/co-TF and signaling proteins. Of those, 3,265 were found to be differentially active (BH-adj. p-value<0.05, Fig. 4, Supplemental Fig. 3). Due to the larger number of nominated proteins, GSEA analysis identified additional statistically significant gene sets, compared to kinase-substrate network analysis. These included metabolism, extracellular matrix organization, cell-cell communication, vesicle-mediated transport, muscle contraction and sensory perception (Fig. 3a). Among the most differentially active proteins, several are involved in Wnt signaling (PIAS4), G protein / RAS / GTPase activity (ARHGAP8, RIT1, RAB23), actin filament and cytoskeleton organization (TPM4, MAP2), vesicular transport (CAVIN1), and others (Fig. 4c).

Applied to the HeLa CCL2 vs. Kyoto SEC-SWATH-MS protein correlation profiles, SECAT identified 214 differential proteins by total abundance, 114 by assembled state abundance, and 64 by monomer abundance. At the level of protein interactions we identified 81 proteins by differential complex abundance, and 76 proteins with different interaction ratios (Benjamini-Hochberg [25] adjusted p-value < 0.01; |log2(fold-change) > 1|) (Supplemental Data 1). Using these results for GSEA with top-level Reactome pathways indicates protein network dysregulation on several levels, affecting cell-cell communication, extracellular matrix organization, vesicle-mediated transport, immune system and signal transduction configurations (Fig. 5a, Methods). Since many features reported by SECAT can be highly correlated, e.g. assembled and complex abundance metrics, a simplistic aggregation to the most significant level per protein provides a reduced overview that can be visualized and augmented by context-specific PPI network integration with cellular compartment GO terms analysis (Fig. 5b, Supplemental Data 7, Methods). The two largest clusters of 89 and 74 proteins inferred by the analysis represent cytosolic and mitochondrial ribosomal proteins, respectively, whereas further macromolecular complexes analysis nominated proteins involved in Complex I and respiratory electron transport. These subnetworks represent large heteromeric macromolecular complexes consisting of proteins whose abundance spans a wide range. However, we found that most of these proteins or complexes are not different in abundance or connectivity between the HeLa CCL2 and Kyoto. Some changes affect well-defined protein complexes as whole, e.g. multiple subunits of the EKC/Keops, Mediator or Arp2/3 Complexes are affected in both, abundance or interactor ratio. Conversely, some proteins with detected differences in complex abundance or interactor ratio are part of larger networks, e.g. cell surface, extracellular space or cell-cell junctions, which illustrates the benefits of representing proteins, PPIs and protein complexes as part of protein networks.

**Figure 5.**
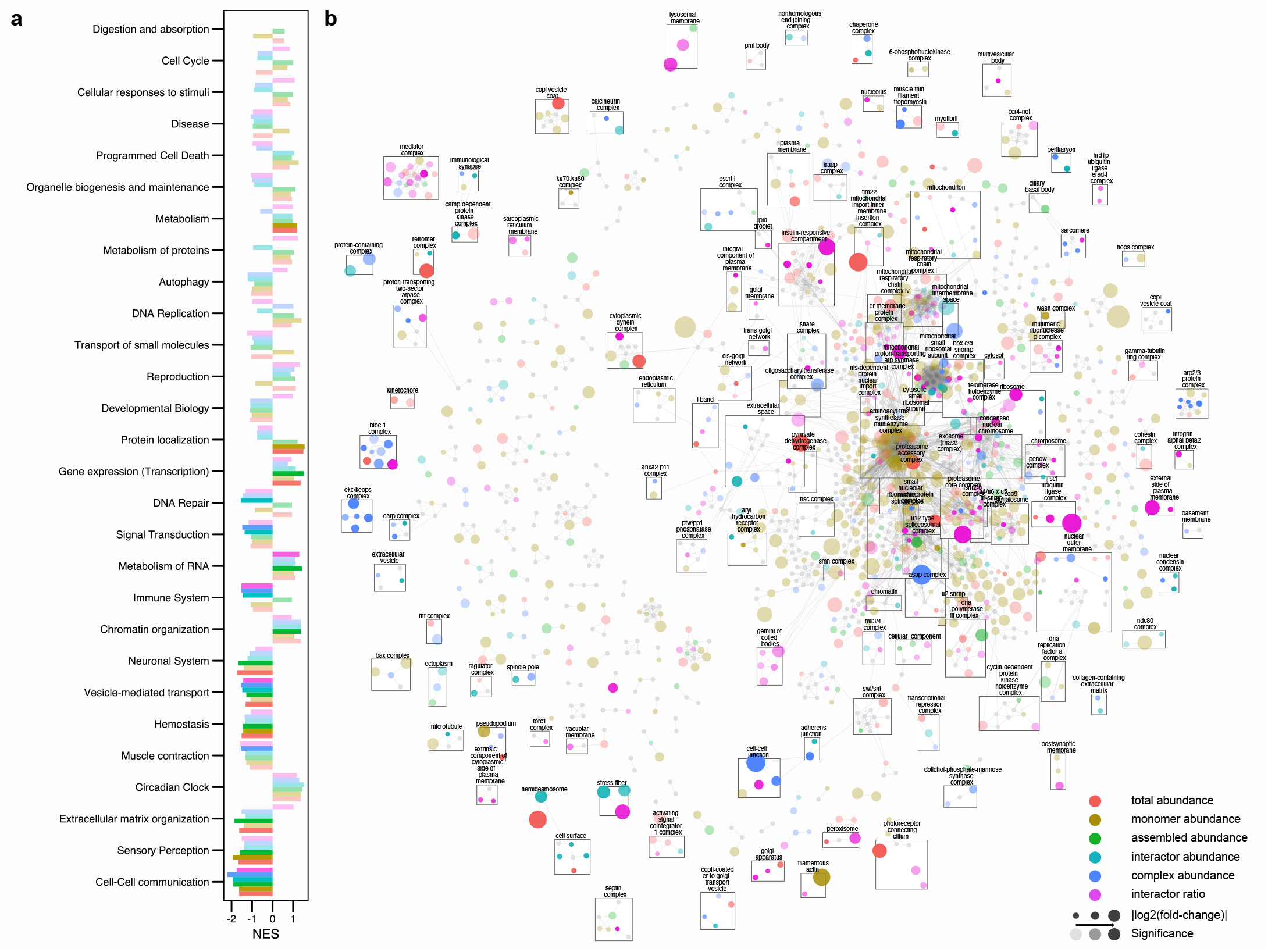
SECAT analysis results. **a)** Gene set enrichment analysis (GSEA) of differential protein complex attributes comparing HeLa Kyoto vs. HeLa CCL2 with top-level Reactome pathways. Opaque categories are significant (BH-adj. p-value < 0.05). **b)** The integrated STRING-based PPI network of the HeLa Kyoto vs. CCL2 data set is depicted with proteins (nodes) and binary PPI (edges) clustered against GO cellular component associations (Methods). Different colors indicate the most significant attribute categories (legend, Methods). Node-size indicates fold-change and opacity indicates statistical significance. Interactive results (Cytoscape) are provided as Supplemental Data 7.

Comparing the sets of differentially active proteins between the three functional levels tested by the algorithms (p. adj < 0.05), illustrates the heterogeneity of protein function, as well as potential confounding technical biases between the different types of analysis (Supplemental Fig. 4). The overlaps range from 1.7% (VESPA-SECAT) to 10.9% (VESPA-VIPER) and 11.1% (VIPER-SECAT). The intersection of VESPA and VIPER highlights kinases/phosphatases which are frequently involved in gene regulation, such as PDGFRA, LIMK1 and MAPK13 (Supplemental Fig. 4a). On the other hand, the intersection between VIPER and SECAT represents proteins with a gene regulatory role, which is frequently mediated by protein-protein interactions, such as the transforming growth factor beta 1 induced transcript 1 (TGFB1I1) (Supplemental Fig. 4b). This confirms the complementarity of these analyses.

Out of the 525 kinases and phosphatases covered by VESPA, only 238 could also be analyzed by SECAT. The majority of these (73.1%) were not differentially active between HeLa Kyoto and CCL2 by VESPA or SECAT analysis (Supplemental Fig. 4c). Since most proteins covered by SECAT are not kinases or phosphatases, we further assessed the relation between differential phosphosite abundance and protein complex dynamics (Supplemental Fig. 4d). Phosphorylation is frequently a core determinant of protein structure and thus can also affect protein complex formation. Several proteins with differential protein complex dynamics, such as the DNA damage-binding protein 2 (DDB2), part of the UV-damaged DNA binding protein complex (UV-DDB), have been found to be differentially phosphorylated. In the case of DDB2, phosphosite S24 has also previously been found to be regulated by PRP4 and it has been suggested to be essential for cancer cell growth [26].

Although SECAT, msVIPER and msVESPA provide functional protein attributes based on different molecular profiles with an only partially overlapping proteome coverage, in general the same perturbed biological processes are identified, suggesting that their complementarity could support integrative analysis based on the inferred PPI and gene regulatory networks.

### Network diffusion elucidates phenotype-associated molecular mechanisms

Genetic or gene regulatory variability often manifests in distinct phenotypes through the proteotype, defined as the contextual state of the proteome encompassing protein abundance, post-translational modifications, and interaction states. Therefore, our goal was to assess whether the context-specific PPI networks and SECAT-informed quantitative features at the different levels of information provided by SECAT could be used to elucidate molecular mechanisms differentiating CCL2 and Kyoto HeLa cells. To identify the effect of upstream signaling or gene regulatory activity and protein complex state changes on downstream layers, and to identify differential network modules between the two subtypes, we used TieDIE [27, 28], a network diffusion approach. As upstream and downstream inputs, we used the top 50 proteins that showed a difference between CCL2 and Kyoto cells based on each data modality, as connected by the context-specific gene regulatory and PPI networks generated above (Methods). Affected pathways between different layers were then identified by GSEA and the first two levels of the Reactome pathway taxonomy were used to summarize the upstream and downstream dependencies (Fig. 6, Supplemental Fig. 5-12, Supplemental Data 8, Methods).

**Figure 6.**
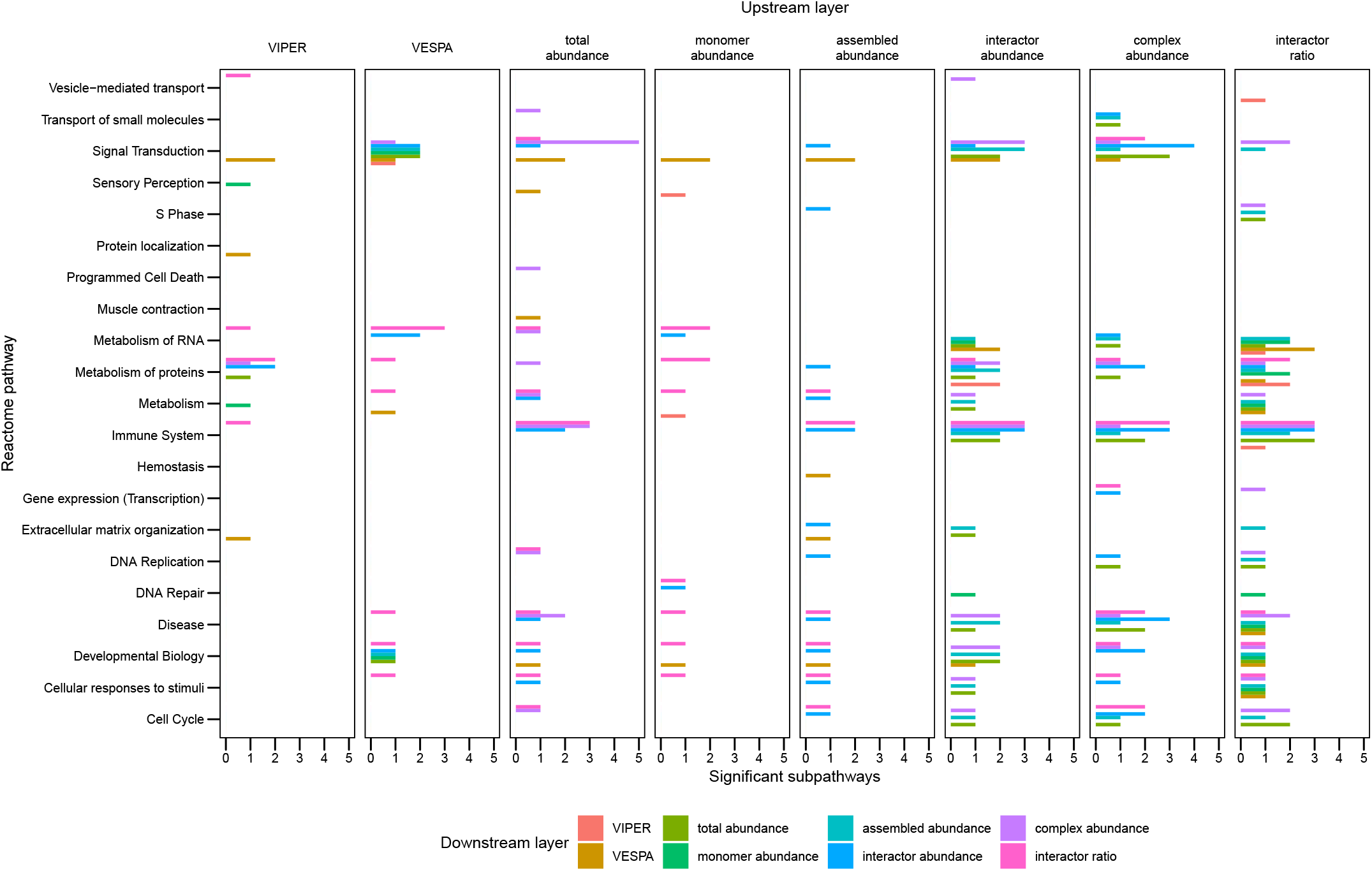
Summary of TieDIE network diffusion analysis results. Horizontal categories indicate upstream layer, the barplot colors (legend) indicate downstream layer categories. The results are split according to top-level Reactome pathways and the number of significant (BH-adj. p-value < 0.05) subpathways. Supplemental Fig. 6-13 and Supplemental Data 8 provide the detailed results and underlying data points.

We used this pathway activation map as guideline to select the following examples to discuss interactions between different data modalities.

#### Effects of kinase activity on protein complexes

Among the most differentially active signaling proteins—as assessed by VESPA—the analysis nominated the LIM domain kinase 1 (LIMK1), showing significantly lower activity in Kyoto compared to CCL2 cells (Fig. 4). LIMK1 was found to be a critical regulator of cytoskeleton dynamics, controlling matrix protein degradation and invadopodia [29]. As part of the “Signaling by Rho GTPases, Miro GTPases and RHOBTB3” pathway (Fig. 6, Supplemental Fig. 5), network diffusion analysis identified Filamin-A (FLNA) and Tropomyosin alpha-3 chain (TPM3) to be affected at the level of protein complex abundance (Fig. 7a), in line with potentially reconfigured cytoskeleton structures and the previously observed lower *Salmonella* typhimurium infection susceptibility of HeLa Kyoto [12].

**Figure 7.**
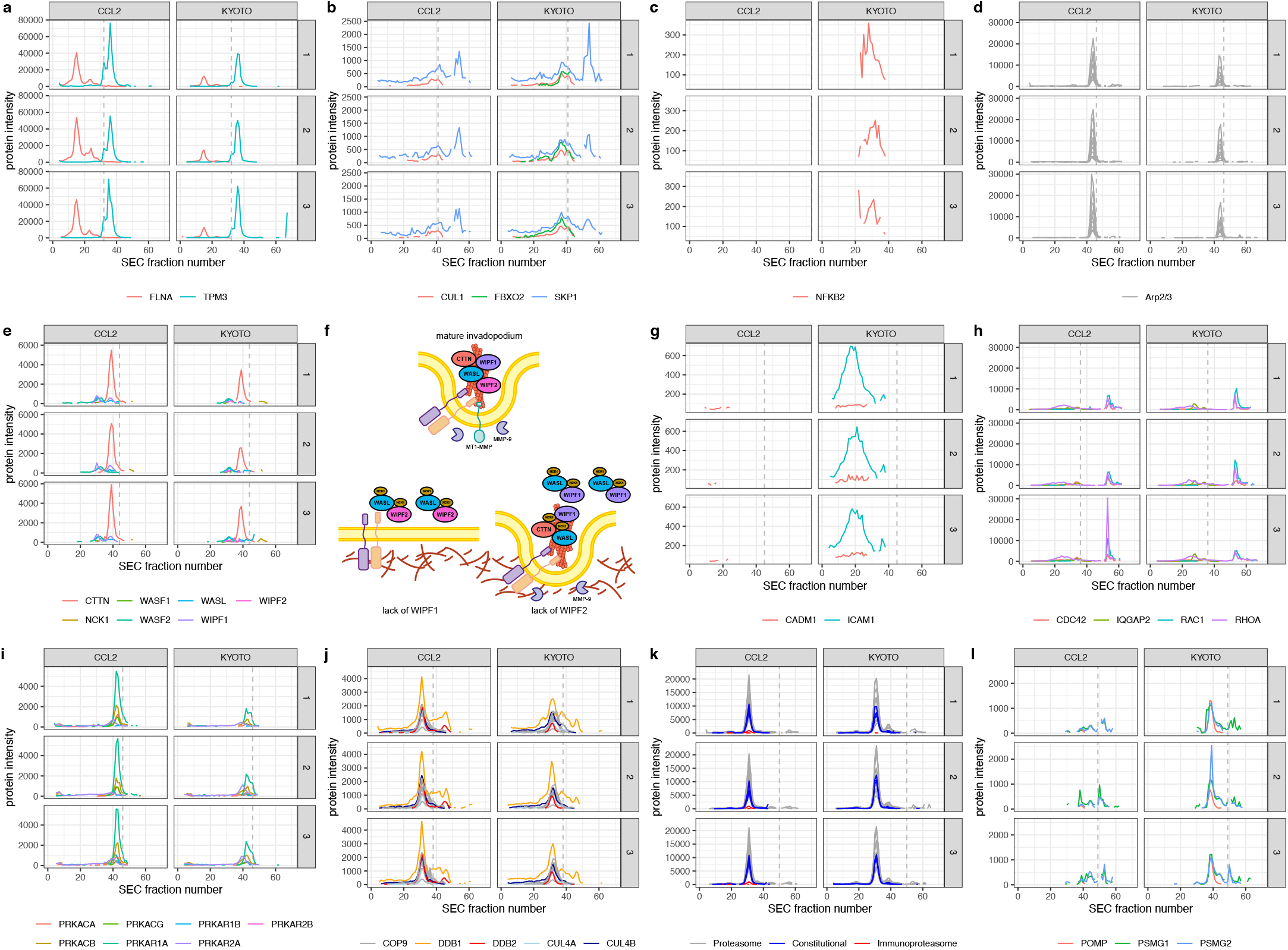
Protein-level SEC-SWATH profiles of HeLa CCL2 and Kyoto cell lines. Lines indicate different protein subunits, whereas the dashed line indicates the highest monomer threshold per group. **a)** Protein complex abundance changes involving Filamin-1 (FLNA) and the Tropomyosin alpha-3 chain (TPM3) between the two cell lines. **b)** Differential configuration of the SKP1-CUL1-F-box (SCF) complex with the FBXO2 substrate recognition subunit. **c)** NF*κ*B is only detectable in HeLa Kyoto cells. **d)** Differential abundance of the actin-related protein complex 2/3 subunits. **e)** Wiskott-Aldrich Syndrome family proteins and their involvement in different subcomplexes. **f)** The molecular mechanism underlying WASP-dependent ECM degradation. Adapted from [40]. **g)** Differential abundance of ECM proteins. **h)** Rho-GTPase activity mediated by IQGAP2 scaffolding. **i)** Differential protein kinase A complex abundance. **j)** Differential complex stoichiometry of COP9 Signalosome-CUL4-DDB1/2 supercomplexes. **k)** Constitutional- and Immunoproteasome abundance differences. **l)** Proteasome maturation protein and Proteasome assembly chaperone expression.

#### Effects of transcription factor activity on protein complexes

Connecting VIPER gene regulatory activity with its downstream regulated targets nominated protein complexes involved in extracellular matrix organization and in other signal transduction pathways as especially affected (Fig. 6, Supplemental Fig. 6). For instance, the Ras-related protein Rab-31 (RAB31), Tumor necrosis factor-inducible gene 6 protein (TNFAIP6) and Integrin alpha-V (ITGAV) have all significantly lower activity in Kyoto vs. CCL2 cells. Moreover, network diffusion analysis identified the S-phase kinase associated protein 1 (SKP1)—part of the SKP1-CUL1-F-box (SCF) ubiquitin ligase complex—to be implicated via the innate immune system and associated processes [30]. The F-box only protein 2 (FBXO2)—one of several substrate recognition component of this complex—was only detected as differentially active in Kyoto cells, suggesting reconfigured mediation of ubiquitin-based degradation [30] (Fig. 7b). Interestingly, the master regulator NF*κ*B (NFKB2) was only detectable as differentially active in Kyoto cells. This represents an established recognition substrate for a different F-box protein, the F-box/WD repeat-containing protein 1A (BTRC) [30], which is not covered by our data set, suggesting that SCF complex the reconfiguration may affect NF*κ*B protein abundance (Fig. 7c).

#### Effects of monomeric proteins on protein complexes

Alterations in upstream monomer abundance primarily affect downstream protein complexes, yet only few other components in our analysis (Fig. 6, Supplemental Fig. 8). The rationale could be that—among the monomers with greater differential activity between the HeLa lines—signaling proteins, receptors and regulators play a critical role in affecting downstream pathways. For example, the Insulin-like growth factor-binding protein 7 (IGFBP7) [31] stimulates cell adhesion and is only detectable in CCL2 cells (Supplemental Fig. 13a). Similar examples include Profilin-2 (PFN2), Calponin-3 (CNN3) and F-actin-monooxygenase (MICAL1), which mediate actin binding and cytoskeleton organization, affecting macromolecular protein complexes (Supplemental Fig. 13a).

Dysregulated cell-cell communication mechanisms represent an important component of carcinogenesis and metastasis, affecting the local tumor microenvironment and cell regulatory processes [32, 33]. In our previous multi-omics study [12], we found relevant differences in CCL2 vs. Kyoto cells, affecting cell doubling time, texture contrast of actin structures, response to perturbation with the tumor suppressor miRNA Let7, and *Salmonella* Typhimurium infection susceptibility. These phenotypes are connected to proteotypic differences in HeLa cells, such as the previously reported differential abundance of Actin-related protein 2/3 (Arp2/3) complex subunits. Arp2/3 has an established role in bacterial internalization by host cells through initiation of membrane ruffles [34–36], also referred to as invadopodia. Consistently, SECAT identified the subunits of Arp2/3 to be differentially abundant between the two cell lines (Fig. 7d).

The subunits of the heteromeric complex are exclusively present in complex-bound conformation, indicating synchronized co-regulation. Arp2/3 is more abundant in HeLa CCL2 and similarly, other actin or cytoskeleton-related protein levels are more abundant or only detectable in this cell line (Supplemental Fig. 14b).

#### Protein complex dynamics

The Arp2/3 complex itself is activated by the Wiskott-Aldrich Syndrome family proteins that include WASP, N-WASP (WASL) and WAVE1/2 (WASF1/2) [37]. While the abundance levels and interactions between WASL and WASF1/2 are not substantially different between the two cell lines, the WASP/N-WASP interacting proteins 1 & 2 (WIPF1/2) are different in abundance and interactions. WIPF1 is more abundant in HeLa CCL2, whereas WIPF2 is only detectable in HeLa Kyoto cell lines (Fig. 7e). The overexpression of WIPF1 has been linked to increased actin polymerization mediated by the WASL-WIPF1 complex [38, 39], which together with cortactin (CTTN) leads to invadopodium initiation and assembly. WIPF2 is further required to give rise to a mature invadopodium that can degrade the extracellular matrix via active matrix metalloproteinases [40]. However, the ratio between WIPF1 and WIPF2 is critical for invadopodium initiation: higher levels of WIPF2 and lower levels of WIPF1 are ascribed to prohibit this step [40] (Fig. 7f). The high ratio of WIPF2 vs. WIPF1 in HeLa Kyoto thus suggests an immature status of the invadopodia which is further supported by the extracellular-matrix protein measurements, including the intercellular adhesion molecule 1 (ICAM1) and the cell adhesion molecule 1 (CADM1) which are less abundant in HeLa CCL2 due to their brake-down by active matrix metalloproteinases [40], but not in HeLa Kyoto (Fig. 7g). Cancer cells frequently have modified surface markers (e.g. antigens, cell adhesion proteins) parts of a crucial process to evade the immune system. They are also differentially configured in HeLa Kyoto vs. HeLa CCL2 cell lines (Supplemental Fig. 13c).

Upstream regulators of WASP family proteins are primarily part of the Rho GTPase protein family. This includes RhoA, Rac1 and CDC42 [37]. While we found these proteins to not be differently abundant between the two cell lines, SECAT identified the Ras GTPase-activating-like protein IQGAP2 in complex with CDC42 to be differently abundant between cell lines (Fig. 7h). Although Rac1 is also a known interactor of IQGAP2, its low measured abundance in complex-bound conformation limits assessment of its PPI. IQGAP2 in complex with CDC42 is more abundant in the HeLa CCL2 cell line, which is in line with its purposed scaffolding function to establish CDC42 and Rac1 activation by inducing a conformational change which mediates GTP binding [41]. Rho GTPases themselves are targeted by upstream kinases, including Src, PKC and PKA [37]. In addition to higher monomeric subunit levels of Src and PKC isomers in HeLa CCL2, SECAT found several proteins part of the protein kinase A complex to be more abundant in HeLa CCL2 (Fig. 7i).

The structural rearrangements of invadopodia formation require elaborate logistics to transport the products from the endoplasmic reticulum via membrane vesicles to their destination. This process is primarily orchestrated by a network of proteins belonging to the Rab GTPase superfamily [42]. Among the 74 superfamily members, 44 are covered by our data set and of those, 7 are differentially abundant between the cell lines (Supplemental Fig. 13d).

It was recently found that *Salmonella* Typhimurium produces the cysteine protease GtgE, which exclusively targets the inactivated form of Rab32, rendering it permanently inactive and thus preventing cells from delivering antimicrobial factors to limit infection [43]. Rab32 is lower abundant as has lower protein gene regulatory activity in HeLa Kyoto, which might indicate that the vesicular transport system is less adapted to prevent bacterial infections.

An essential part of the cellular immune system is protein degradation. The COP9 signalosome (CSN) is a heteromeric protein complex primarily involved in post-translational protein regulation of Cullin-RING ubiquitin ligases (CRLs), however, recent studies also implicated involvement in cancer-associated mechanisms [44]. We found CSN subunit levels to be similar between the two cell lines, but we found differential stoichiometries of associated complex-bound DNA-damage binding proteins 1/2 (DDB1/2) (Fig. 7j). In response to DNA damage, it was proposed that CSN forms complexes with CRLs, acting on chromatin to promote DNA repair and checkpoint control [45]. The observed differential DDB1/2 complex stoichiometries might originate from subcomplexes which are not separable by SEC. It was proposed that different supercomplexes binding DDB1/2 have specific roles and mechanisms in tumor checkpoint control, however the exact mechanisms are not yet understood [45]. A similar adaptation can be observed in the related ubiquitin proteasome system: In HeLa Kyoto, only subunits of the heteromeric constitutive proteasome complex are observed above the limit of detection (Fig. 7k). In HeLa CCL2 however, the observed peak can be attributed to both constitutive and immunoproteasome, a variant of the proteasome with increased activity during inflammation, also supporting antigen representation of different peptide fragments by MHC class I molecules [46]. Interestingly, the proteasome assembly chaperones 1/2 and the proteasome maturation protein are higher abundant in HeLa Kyoto, which might indicate a preferred association with the constitutional proteasome (Fig. 7l).

In summary, these findings support the differential and coordinated modulation of the cellular immune system, cytoskeleton, cell surface and extracellular matrix in HeLa Kyoto vs. CCL2 starting from signaling pathways to vesicle-mediated transport of structural proteins and the ECM by covering the involved protein complexes.

## Discussion

The measurement of comprehensive, multi-omic (genome, transcriptome, proteome, metabolome) molecular profiles across large cohorts has become accessible for many study types due to recent technological advances. Even though proteins are more closely associated with biological function than genes, transcripts or metabolites, their measured molecular abundance frequently does not directly correlate with the protein’s activity thus impeding the association of protein abundance with individual phenotypes. This divergence of abundance and function is caused by several mechanisms acting on proteins, including post-translational modifications, three-dimensional structure, association with other biomolecules to form complexes and combinations thereof, that modulate the inherent activities and functions of proteins [47]. Here, we suggest that inference of functionally relevant protein attributes, specifically signaling or gene regulatory activity and protein complex state, can be more informative than mere protein abundance and provide direct mechanistic insights associated with phenotypes.

To test this hypothesis, we conducted a study that compared at a high level of detail two phenotypically diverse instances of a panel of HeLa cells that have diversified by genetic drift. For this purpose, we acquired protein correlation profiles of the cell lines in exponential growth and differentially quantified the protein network state by SECAT, msVESPA, and msVIPER at the level of PPI networks, signaling and gene regulation levels, respectively. We could then use this representation of molecular interactions by network diffusion analysis to assess the mechanistic effects of signaling activity, gene regulatory activity and protein complex states on each other. Further, quantitative assessment supported our previously raised [12] hypothesis of “protein complex buffering”, by demonstrating that proteins in assembled configurations are not as directly affected by CNV and mRNA abundance changes as their monomeric counterparts.

Our application to the HeLa Kyoto vs. CCL2 cell line comparison further illustrates the potential of protein correlation profiling for mechanistic investigations, requiring only three replicates per condition to identify many differentially connected modules. In addition to distinct, unambiguous protein complexes such as the proteasome (Fig. 7k), differential protein complex engagement using non-Gaussian protein complex profiles (e.g. WASP associated proteins, Fig. 7e) can be assessed to permit an unbiased overview of functional proteome changes between experimental conditions. This enabled us to extend our previous study on the differences of these two cell line types and identified adaptations regarding cell surface and immune system, supporting previous findings on the involved molecular mechanisms.

While the present work employs HeLa cells as a model, their extensive characterization across genomic, transcriptomic, proteomic, and phenotypic layers [12] makes this system uniquely well-suited for the further interrogation of multifaceted protein attributes such as complex assembly states and regulatory network activities beyond protein expression levels. Importantly, our integrative framework demonstrates how such multifaceted attributes, derived from both measured complex dynamics and inferred activities, can more effectively connect molecular perturbations with emergent phenotypic outcomes. Beyond HeLa, we envision our approach presented here as a general paradigm for studying proteins as adaptive agents whose diverse attributes jointly shape cellular states within complex biological systems.

In conclusion, the study shows that new methodologies aiming at the analysis of contextual network analysis provide informative insights into the molecular mechanisms associated with or causative of phenotypes and covers the state of transcriptional programming, characterized by protein gene regulatory activity, as well as the involvement of proteins in signaling networks or complexes, characterized by their substrates or complex stoichiometries. Although these networks operate differently with only a partial overlap, the transformation of functional attributes to individual protein subunits makes data integration and interpretation straight-forward and can provide complementary and independent insights with relevance for the understanding of molecular mechanisms.

## Data availability

The HeLa Kyoto vs. CCL2 SEC-SWATH-MS mass spectrometry proteomics data and primary data analysis results have been deposited to the ProteomeXchange Consortium via the PRIDE [48] partner repository (https://www.ebi.ac.uk/pride/archive/projects/PXD067680) with the data set identifier PXD067680.

## Code availability

All code and supplemental datasets are available on Zenodo (https://zenodo.org) with the following DOI: https://dx.doi.org/10.5281/zenodo.16985688

## Acknowledgments

This study was supported by NCI U54CA274506 (Center for Cancer Systems Therapeutics, CaST), a supplemental grant to NCI U54 CA209997 (Cancer Systems Biology Consortium), the NCI Office of Cancer Target Discovery and Development (CTD2) award U01CA272610, and the NIH Shared Instrumentation Grants S10 OD012351 and S10 OD021764 all to A.C. G.R. was supported by grants P2EZP3_175127 and P400PB_183933 from the Swiss National Science Foundation. Y.L. was supported by the National Institute of General Medical Sciences (NIGMS), NIH through grant R01GM137031 and R35GM158073.

## Author contributions

- George Rosenberger: Conceptualization, Methodology, Software, Formal analysis, Investigation, Writing - Original Draft, Visualization
- Peng Xue: Methodology, Investigation, Data Curation
- Isabell Bludau: Methodology
- Ben Collins: Methodology
- Claudia Martelli: Methodology
- Evan Williams: Methodology
- Andrea Califano: Methodology, Formal analysis, Writing - Review & Editing
- Yansheng Liu: Conceptualization, Investigation, Writing - Review & Editing, Supervision, Funding Acquisition
- Ruedi Aebersold: Conceptualization, Writing - Original Draft, Supervision, Funding Acquisition

## Methods

### Protein complex dynamics of HeLa Kyoto vs. CCL2 cell lines

#### Sample preparation

HeLa CCL2 (HeLa 14) and Kyoto (HeLa 8) cell lines were obtained and cultured as described in our previous study [12].

#### SEC protein complex fractionation

Protein complexes fractionation was performed as previously described [10]. HeLa CCL2 cells and HeLa Kyoto cells were thawed and lysed in mild conditions by homogenization with a lysis buffer composed of 0.5% NP-40 detergent and protease and phosphatase inhibitors (50 mM HEPES pH 7.5, 150 mM NaCl, 0.5% NP-40). Cell debris and membranes were removed by 15 minutes of ultracentrifugation (50,000×g, 4 °C) and the detergent was removed by 30 kDa molecular weight cut-off membrane and exchanged with the SEC buffer (50 mM HEPES pH 7.5, 150 mM NaCl). After 5 min of centrifugation at 16,900 ×g at 4 °C, the supernatant was directly injected to a SRT-C-SEC 500 column (dimensions 300×21.2 mm, pore size 500 Å, particle size 5 µm, Sepax-Tech, DE, USA). Per SEC run, 2.6 mg of native proteome extract (estimated by Pierce™ BCA Protein Assay Kit, Thermo Scientific) was injected and fractionated at 2 ml/min flow rate on ice (4 °C), collecting 90 fractions at 0.4 min per fraction from 18 to 54 min post-injection. According to the protein elution window, fractions 1-70 (18 – 46 min elution time) were collected for proteomic analysis. The calibration curve for SEC fractionation was obtained by measuring a protein standard mix (Column Performance Check Standard, Aqueous SEC 1, AL0-3042, Phenomenex, CA, USA) before each sample.

#### Sample preparation for mass spectrometry

Sample processing for bottom-up analysis of SEC fractions was performed on a 96-well plate MWCO filters (AcroPep Advance Filter Plates for Ultrafiltration 1mL Omega 10K MWCO; Pall Corporation, USA) [49]. Prior to usage, the filters were washed twice with 200 µl of water that was successively removed by centrifugation at 1800 g for 30 min. 70 fractions for each sample were loaded and concentrated on the filters through centrifugation, until complete removal of the SEC buffer. Protein denaturation and reduction was obtained after incubating the samples at 37 °C for 30 min with 5 mM of TCEP in 8M Urea/20 mM AMBIC (pH 8.8). Alkylation of cysteine residues was performed by adding a final concentration of 50 mM IAA/20 mM AMBIC and incubating in the dark at room temperature for 1 h. After the reaction, the plates were centrifuged to remove Urea buffer and washed for three times with 20 mM AMBIC. Protein digestion was carried out at 37 °C for 16 h, adding to each fraction 1 µg of trypsin (Promega, Switzerland) and 0.3 µg of Lysyl Endopeptidase (Mass Spectrometry grade, FUJIFILM Wako Pure Chemical Industries, Japan). The resulting peptides were collected by centrifugation and further washing of the filters with 20 mM AMBIC.

#### Mass spectrometry

LC-SWATH-MS analysis of the peptide fractions was performed on Evosep One system (Evosep Biosystems, Denmark) [50] coupled to a SCIEX TripleTOF 6600 instrument (SCIEX, MA, USA) equipped with a NanoSpray III ion source (SCIEX). Due to the heterogeneity of the SEC proteins fraction, we normalized the injections according to the fraction volume and to the amount of protein fractionated, corresponding to a total amount of 325 µg of proteins.

#### Sample loading

The samples are loaded in Evotips (Evosep Biosystems, Denmark), after resuspension in solvent A (0.1% FA water solution; Fisher Scientific AG, Switzerland) and the addition of iRT-Kit peptides (Biognosys AG, Switzerland). Prior to loading, the C18 stage tips (Evotips) were soaked with 100 µl of 2-propanol during the activation and the conditioning steps. The activation step consisted in the washing with 20 µl of solvent B (0.1 % FA in ACN, Fisher Scientific AG, Switzerland), followed by the conditioning with 20 µl of solvent A. Prior the sample loading step, 10 µl of solvent A was added on top of the tips, ensuring that the tips remained wet during the loading step. For all the steps the tips have been centrifuged for 1 min at a speed of 700 g. The last step (i.e. washing step) was performed using 100 µl of solvent A, and the loaded tips were added with 200 µl of solvent A for preserving the samples soaked during the entire injection of the batch.

#### Evosep-ESI-DIA-SWATH

The separation of peptides was performed selecting the “60 samples per day” method, consisting in 24 minutes of total cycle time, for 21 minutes of gradient length, 3 minutes of overhead time at a flow rate of 1 µl/min. A partial gradient is applied (0-35% solvent B) in order to elute the peptides from the Evotips by a couple of low-pressure pumps. The peptides were then pushed in a C-18 nanoConnect LC column (8 cm column, ID 100 µm packed with 3 µm Reprocil, PepSep, Denmark) using a high pressure pump and solvent A [50]. The ESI coupling was obtained using a Nano Source Emitter Stainless Steel (Fisher Scientific AG). The ESI tuning parameters were the following: spray voltage, 2800 V; ion source gas flow (GS1), 16; curtain gas flow (CUR), 35; interface heater temperature (IHT), 100°C and declustering potential, 100. The Evosep system was controlled by the Axel Semrau Chronos software (Axel Semrau GmbH, Germany), while the mass spectrometer acquisition software was Analyst TF 1.7.1 (Sciex).

Data-independent acquisition mass spectrometry was performed for the quantitative analysis of the 420 SEC fractions (70 fractions per sample) obtained from the 6 SEC experiment. SWATH scans were performed using an updated scheme of 64 variably sized precursor co-isolation windows [51], covering similar precursor densities (in terms of number and intensity) within all SWATH windows. The SWATH windows covered the precursors ions in the range of 350-1500 m/z and 350-1500 in the MS2 SWATH scans, the accumulation time was 100 ms for the MS1 and 20 ms for each SWATH window, resulting in a cycle time of 1.38 s. For fragmentation, a rolling collisional energy with a collisional energy spread of 15 eV was applied.

#### Primary data analysis

The mass spectrometric data has been analyzed with Spectronaut [52] (Biognosys AG, Schlieren, Switzerland; software revision 12.0.20491.11.25470) and the software-provided version of our previously published combined human spectral library [53]. The vendor-suggested parameters have been used, except the normalization of peptide abundances, which has been deactivated. All mass spectrometry raw data, the exact used set of parameters and the processed primary data analysis results are available on the ProteomeXchange repository.

#### Data analysis

SECAT [11] (version 1.0.6), PyProphet [54, 55] (version 2.1.5) and VIPER [4] (version 1.20) were used for all data analyses with STRING (version 11.0 [18]), CORUM (version 3.0 [17]) and PrePPI (version 2016 [56]) and default parameters if not otherwise specified. The raw profiles were normalized within SECAT as described previously [11] (Supplemental Fig. 15-16). Semi-supervised learning was conducted using CORUM as positive network and CORUM-inverted as negative network. All input data and parameters are provided on the Zenodo repository. CORUM-inverted was generated as described before [11] by using the inverted set of PPI (i.e. all possible PPI that are not covered by CORUM) and removing all PPI in this set covered by STRING [18], IID [57], PrePPI or BioPlex [58].

The gene set enrichment analysis in Fig. 5a was generated using fgsea (version 1.18.0) [20], applied to the signed SECAT scores on all levels ((sign of log2(fold-change)*(-log10(BH-adj. p-values))). The Reactome Pathway Database [59] (version 77; only top-levels) was used with default parameters. Opacity in the figure indicates significance (BH-adjusted p-value < 0.05).

To annotate and visualize differential proteins in Fig. 5b between the HeLa CCL2 and HeLa Kyoto cell lines identified by SECAT, we used Cytoscape [60] (version 3.8.0). GO Cellular Components annotation (version 20200616) was used to cluster PPI using the Cytoscape App AutoAnnotate [61] with default parameters and a maximum cluster size (“Max words per label”) of 1. Clusters were arranged according to the CoSE layout.

Supplemental Fig. 14-15 were generated as described previously [11].

### Quantitative effects of copy number variation and mRNA abundance on protein complex assembly states

The multi-omic data has been obtained from the original publication [12]. The SECAT quantitative values have been filtered to ensure accurate measurements between replicates: The standard deviation was computed for the three replicates of both biological conditions separately and aggregated to the maximum of the two conditions. Only proteins with a score lower than the 75% quantile of all scores on total-level were compared.

Spearman’s *ρ* correlation coefficients are reported. Statistical tests between the correlation coefficients were conducted as proposed using Fisher’s transformation [62].

### Association of protein abundance and protein complex/network dysregulation with phenotypic variability

#### Data preprocessing

Whole proteome quantitative profiles have been obtained from our previous study [12]. OpenSWATH peak-group level results were imported and aggregated to protein-level using the R-package “iq” [63] (version 1.9). Protein abundances were then z-score transformed over all samples. For recursive feature elimination of protein abundance features, only “central culture” samples were used. For the DeMAND analysis, missing values were imputed bi a row-wise minimum, including jitter.

#### Inference of complex and network dysregulation

To infer metrics for dysregulation of the protein network state from whole proteome samples, we used the *Salmonella* Typhimurium infection proteomic data from our original study [12], where for each cell line the proteome was measured in duplicates as part of a control experiment and after textitSalmonella Typhimurium infection. Based on this quantitative peptide matrix, we inferred protein abundance and generated context-specific PPI networks based on the generalized CORUM [17] (protein complexes) and STRING [18] (protein network) reference databases using the ARACNe [64, 65] algorithm applied to the protein abundance quantitative matrix. To assess complex- or network-level dysregulation of PPIs, we used the DeMAND [19] algorithm in combination with the protein-level quantitative matrix and the context-specific complex and network interactions from ARACNe.

##### PPI network reconstruction

ARACNe (Git revision: 4ae398c) was used using the inferred protein abundances to generate separate context-specific subsets of CORUM [17] (version 3.0) and STRING [18] (version 11.0). The bidirectional PPI networks were used to constrain the query space and ARACNe was used with default parameters and 200 bootstrap runs. All PPIs with a BH-adj. p-value < 0.05 were used for the consecutive steps.

##### DeMAND analysis

DeMAND (version 1.22.0) was used with the two subset PPI networks separately and the inferred protein abundances to estimate PPI and protein network state dysregulation for each cell line. For this purpose, Let7-perturbed and control samples of a cell line were selected as foreground and the control samples of all cell lines were used as background profiles. DeMAND was run with default parameters.

#### Recursive feature elimination to identify the most predictive features

Features from all data modalities were constraint to the least common denominator: Only protein abundances covered by the STRING-DeMAND analysis were included.

For each data modality (protein abundance, protein complex dysregulation, protein network dysregulation) and phenotype (cell doubling time, infection rate) separately, recursive feature elimination (RFE) was then conducted using the function “rfe” from the R-package caret [66] (version: 6.0-86), by Random forest regression with 500 bootstraps and otherwise default parameters. Between 2^0^ – 2^12^ features were selected for estimation of R2 and RMSE metrics.

#### Gene set enrichment analysis

GSEA was conducted using the R-package fgsea [20] (version: 1.14.0) against separate GO databases obtained from MSIGDB [21] (version 7.4) with a minimum gene set size of 5, a positive score type and otherwise default parameters. For each data modality and phenotype, variable importance as reported by RFE was used as scoring metric. Only primary gene sets after pathway collapsing were visualized, which achieved BH-adj. p-value < 0.05 in at least one category in Fig. 2 and Supplemental Fig. 1-2. Fig. 2a depict boxplots with the following parameters: Lower and upper hinges represent the first and third quartiles; the bar represents the median. Lower and upper whisker extend to 1.5 * IQR from the hinge. This represents the default parameters of the function “geom_boxplot” of ggplot2 (3.3.2).

### Differential kinase/phosphatase, transcription factor activities and protein complex dynamics of HeLa Kyoto vs. CCL2 cell lines

#### Differential phosphopeptide abundance analysis

Phosphoproteomic profiles were obtained from our previous study [23]. Phosphopeptides were mapped to separate sites and intensities were log2-transformed. The LIMMA R-package (version 3.52.4) was used to conduct differential abundance analysis on phosphopeptide-level. Multiple-testing corrected p-values were used to generate the correlation plots (Supplemental Fig. 4).

#### Inference of signaling activity

##### Preprocessing

Phosphoproteomic profiles were obtained from our previous study [23]. Phosphopeptides were mapped to separate sites and intensities were log2-transformed. Missing values were imputed row-wise as described previously [7]. All values were rank-normalized [67].

##### Signaling network generation

The dVESPA [7] module (vespa version 1.0.0, vespa.networks version 1.0.0, vespa.db version 1.0.0) was used with default parameters for signaling network generation, as described previously. The preprocessed HeLa phosphoproteomic profiles were used to select the most representative signalons from a collection of 10 CPTAC cancer datasets (COAD [68], OV [69, 70], BRCA [71], UCEC [72], CCRCC [73], LUAD [74], PBT [75], GBM [76], HCC [77], HNSCC [78]). HSM/P1-constrained substrate-level signalons were used for all following steps.

##### Differential Signaling activity inference

msVESPA [7] (vespa version 1.0.0, viper version: 1.26.0) was then used to compare HeLa Kyoto vs. HeLa CCL2 cell lines. msVESPA was configured with 1,000 bootstraps (averaged by mean) and the parametric approach with the meta-signalons generated from above, adaptively restricted to the top 50 substrates. Pleiotropic cross-talk correction was enabled as described previously [7]. Results are reported with BH-adjusted p-values and with mapped identifiers respectively, using Bioconductor/org.Hs.eg.db: 3.13.0.

#### Inference of protein gene regulatory activity

##### Preprocessing

Transcriptomic profiles were obtained from our previous study [12]. Transcript RPKM values were averaged per Entrez identifier, log2-transformed and rank normalized as described previously [67]. Gene regulatory network generation: All TCGA-based ARACNe [64, 65] networks [67] were used to generate an optimal set of regulons by the metaVIPER approach, as described previously [24]

##### Differential TF activity inference

msVIPER [4] (viper version: 1.26.0) was then used with 1,000 bootstraps (averaged by mean) and the parametric approach with the meta-regulon generated above to compare HeLa Kyoto vs. HeLa CCL2 cell lines. Results are reported with BH-adjusted p-values and with Ensembl and UniProt identifiers respectively, mapped using Bioconductor/org.Hs.eg.db: 3.13.0.

#### Gene set enrichment analysis

The gene set enrichment analysis in Fig. 4a was generated using fgsea and default parameters (version 1.18.0) [20], applied to msVESPA and msVIPER NES scores. The Reactome Pathway Database [59] (version 77; only top-levels) was used. Highlighted bars in the figure indicate significance (BH-adjusted p-value < 0.05). Fig. 4b depicts boxplots of leading edge targets with the following parameters: Lower and upper hinges represent the first and third quartiles; the bar represents the median. Lower and upper whisker extend to 1.5 ∗IQR from the hinge. This represents the default parameters of the function “geom_boxplot” of ggplot2 (3.3.5). The GSEA bars represent associated leading edges of the individual genes with the top Reactome pathways. Supplemental Fig. 3 was generated analogously, however with the full Reactome pathway database.

#### Network diffusion elucidates phenotype-associated molecular mechanisms

The context-specific SECAT protein-protein interaction network at 5% global q-value cutoff was used for all analyses, covering 8,978 PPI. To focus on the most confident signaling and gene regulatory interactions, a filtered network was constructed from the optimized VESPA and VIPER signalon/regulon targets after mapping to UniProt identifiers, as described above.

Signaling interactions were selected if both regulators and targets were marked as leading edge by msVESPA and present in either SECAT network or in the msVIPER or msVESPA results, resulting in 8,620 signaling interactions. Gene regulatory interactions were selected if both regulators and targets were marked as leading edge by msVIPER and present in either SECAT network or in the msVIPER or msVESPA results and if the differential regulator activities fulfilled the significance (BH-adjusted p-value < 0.01) and probabilistic weight (l > 0.9) thresholds, resulting in 11,892 gene regulatory interactions. In combination, this provided an integrated network for network diffusion analysis.

To select the top 50 upstream and downstream differential proteins for network diffusion analysis, different schemes were applied: For SECAT “total”, “assembled” and “interactor” abundance levels, significant proteins (BH-adj. p-value < 0.05) were selected according to descending absolute log2(fold-change). For SECAT “interactor”, “complex” abundance and “interactor ratio” levels, proteins were selected according to increasing statistical significance. For msVESPA and msVIPER, proteins were selected according to descending absolute NES.

A custom build of TieDIE [28] (https://github.com/grosenberger/TieDIE/tree/feature/kernel; revision 8a1b583) was applied using the combined network and upstream/downstream inputs described above. First, a kernel was generated for the combined network and then all upstream and downstream input permutations were computed using default parameters. TieDIE heat scores were then used for the GSEA in Fig. 6 and associated data, which was generated using fgsea (Bioconductor/fgsea: 1.18.0) [20]. The Reactome Pathway Database [59] (version 77; filtered to the second level) was used with default paramters. The overview in Fig. 6 indicates significant subpathways for all top-level pathways. Opacity in Supplemental Fig. 5-12 indicates significance (BH-adjusted p-value < 0.05).

Visualization of protein-level SEC-SWATH-MS profiles in Fig. 7 and Supplemental Fig. 13 was conducted using the R-package “ggplot2” by averaging the three most intense peptide precursors per protein.

## Supplemental Figures

**Supplemental Figure 1.**
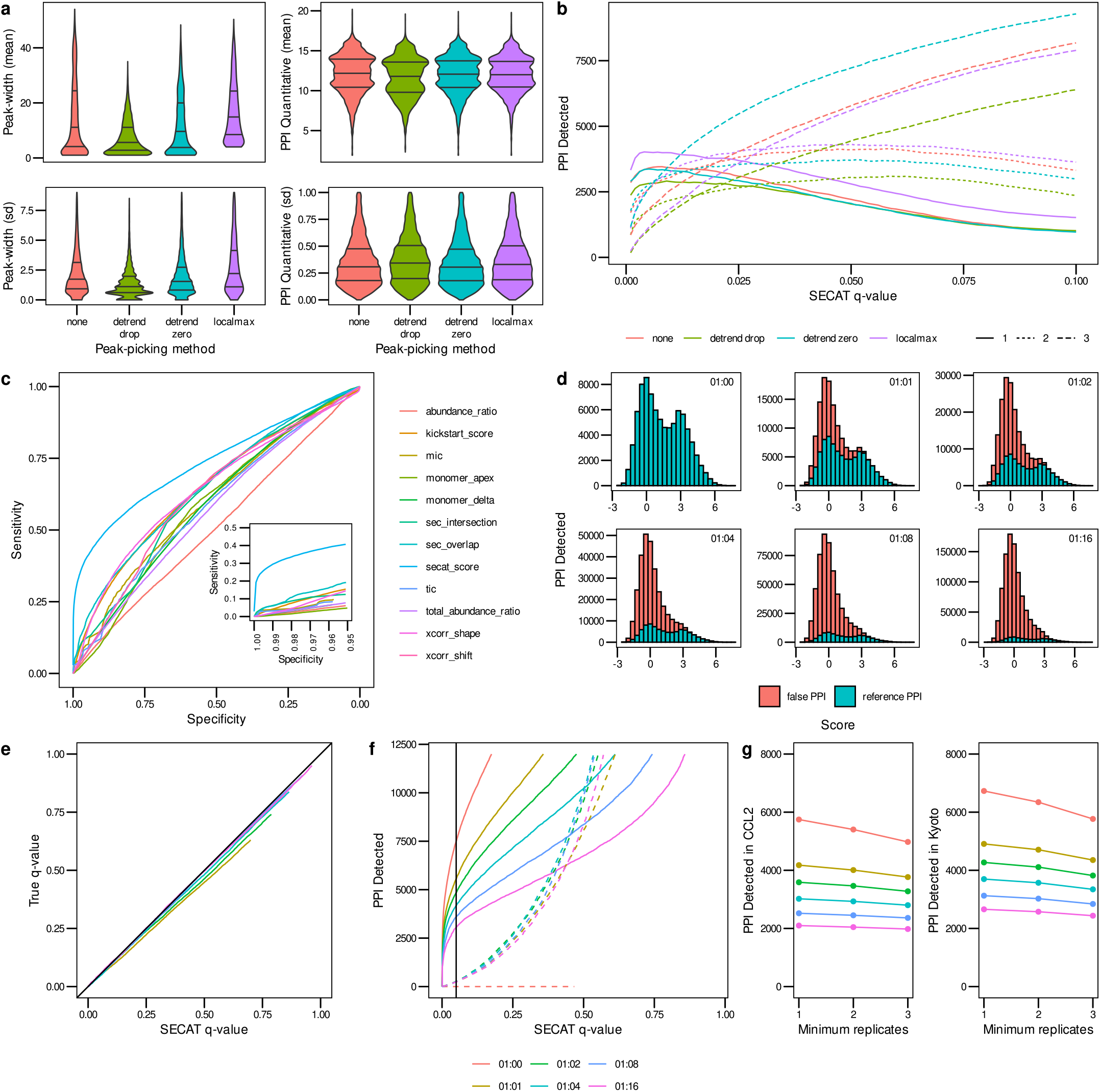
SECAT signal processing and PPI detection performance. **a)** The effect of different peak-picking methods for signal processing on peak-width and PPI quantification within replicates of the same experimental condition is depicted in boxplots (Methods). **b)** Sensitivity of PPI detection vs. SECAT q-value for peak-picking options. The PPI were filtered to a global context q-value<0.05. Colors indicate different peak-picking options; line types indicate number of detections with the three replicates per condition. **c)** The receiver-operating characteristic (ROC) illustrates the sensitivity vs. specificity of the different SECAT subscores, the kickstart score and the integrated SECAT score. **d)** The SECAT score histograms depict the dilution of the reference PPI network (true: CORUM) with false PPI (false: CORUM-inverted) for the benchmark (Methods). **e)** The SECAT estimated q-value is accurate as evaluated against the ground truth for all levels of reference PPI network accuracy. **f)** The sensitivity of PPI detection in dependency of SECAT q-value for different levels of reference PPI network accuracy is depicted. **g)** Consistency of PPI detection between replicates of the same conditions.

**Supplemental Figure 2.**
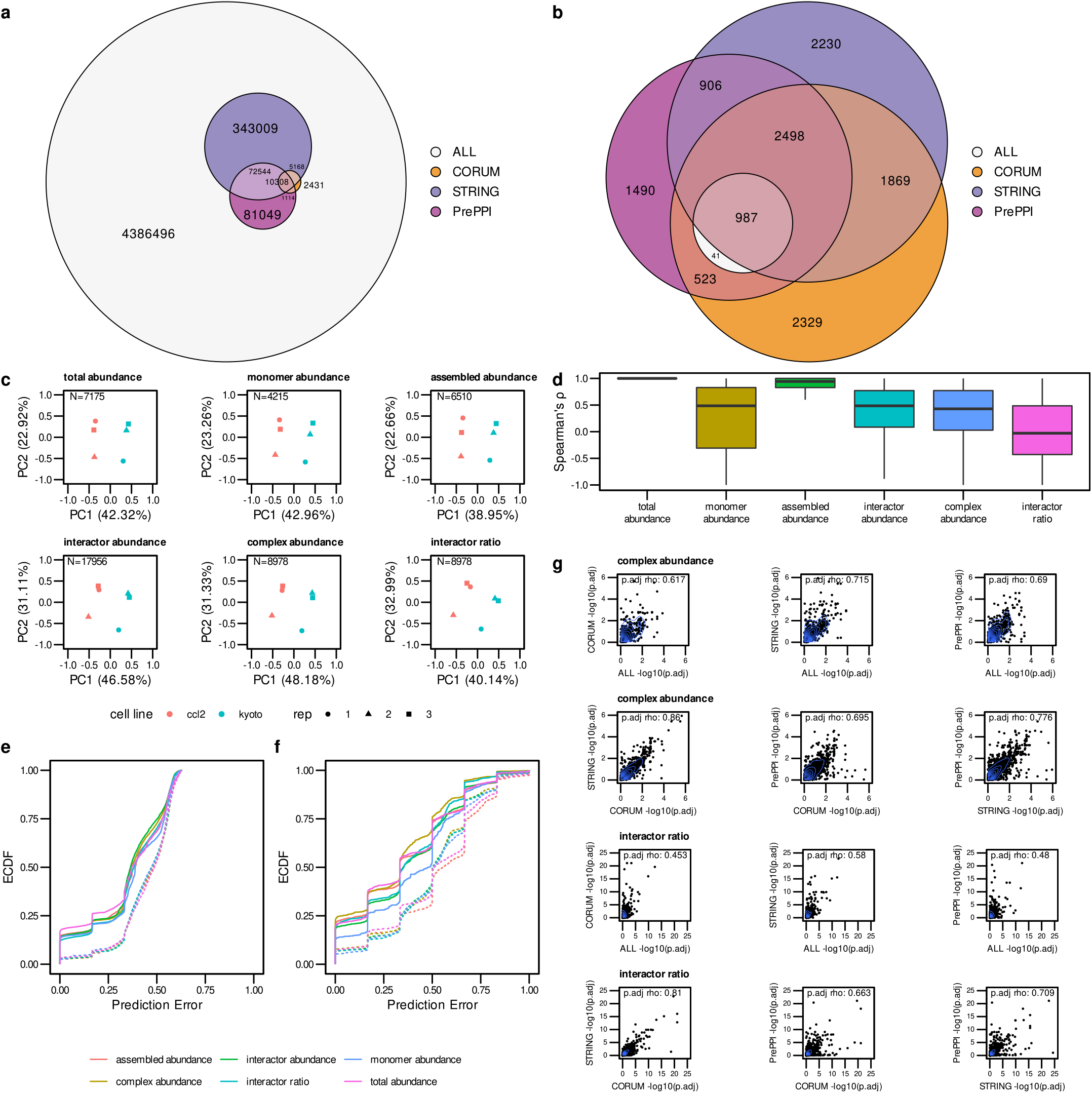
SECAT PPI quantification and Network integration. **a)** The Euler diagram indicates the overlap of STRING, PrePPI and CORUM reference PPI networks resulting in at least a partial overlap of the interactors in any of the replicates. **b)** Overlap of confident (q-value < 0.05) PPI detections after SECAT analysis with different reference PPI networks. **c)** Quantitative SECAT features can be used to separate experimental conditions and replicates. **d)** Spearman’s correlation of features (PPI: summarized by averaging) to total-level features is depicted in individual boxplots. **e)** Predictive error for feature-wise leave-one-out cross-validated logistic regression classification of replicates to biological conditions. **f)** Predictive error for group-wise leave-one-out cross-validated logistic regression classification of replicates to biological conditions. Groups represent several features connected to a core feature based on PPI network. **g)** Correlation of node-level integrated BH-adj. p-values estimated using different reference PPI networks.

**Supplemental Figure 3.**
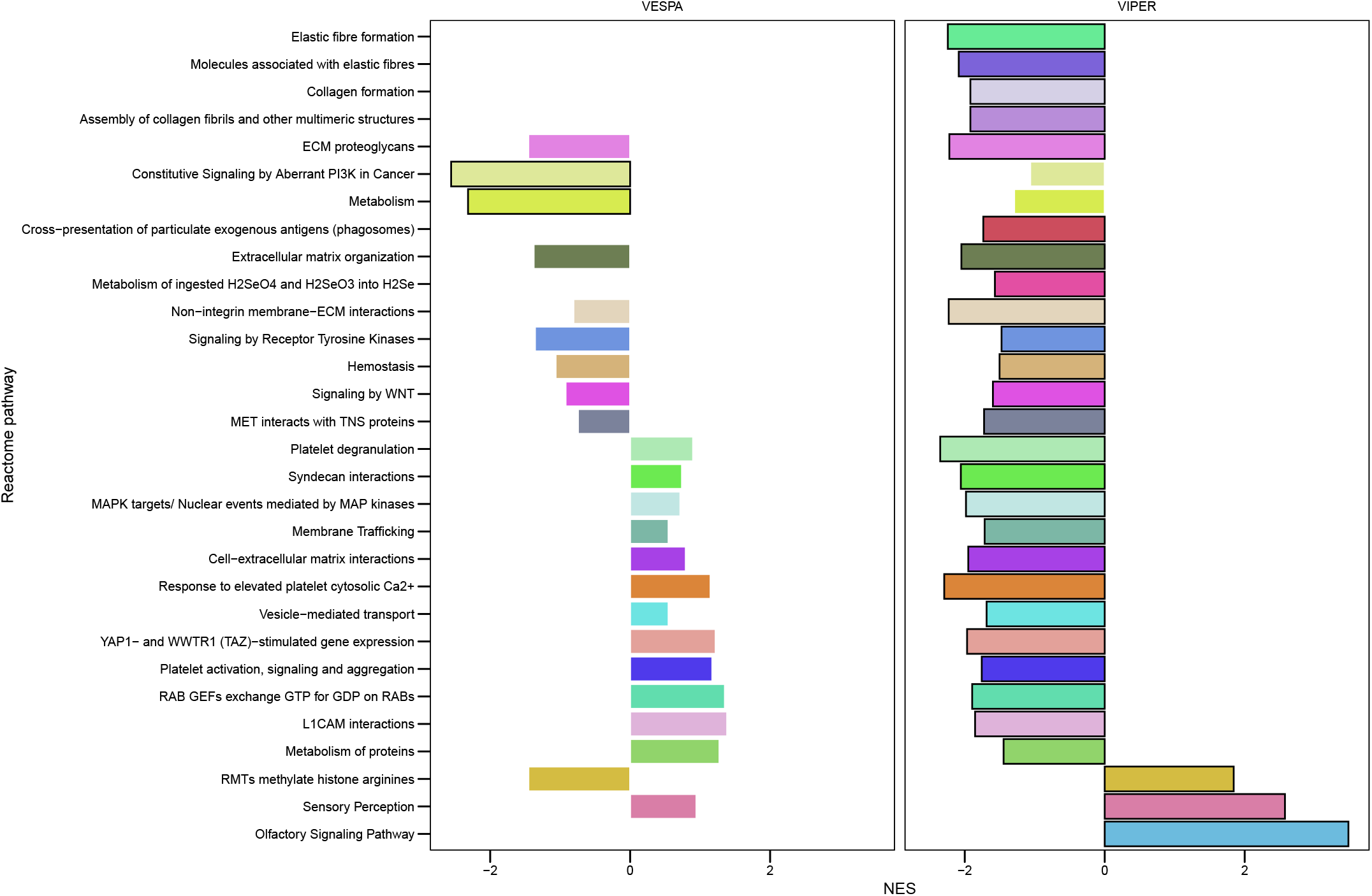
Full Reactome pathway analysis of differential protein regulatory and signaling activities. Gene set enrichment analysis (GSEA) against all Reactome pathways indicates differences between the cell line subtypes. Bold highlighted categories are significant (BH-adj. p-value < 0.05).

**Supplemental Figure 4.**
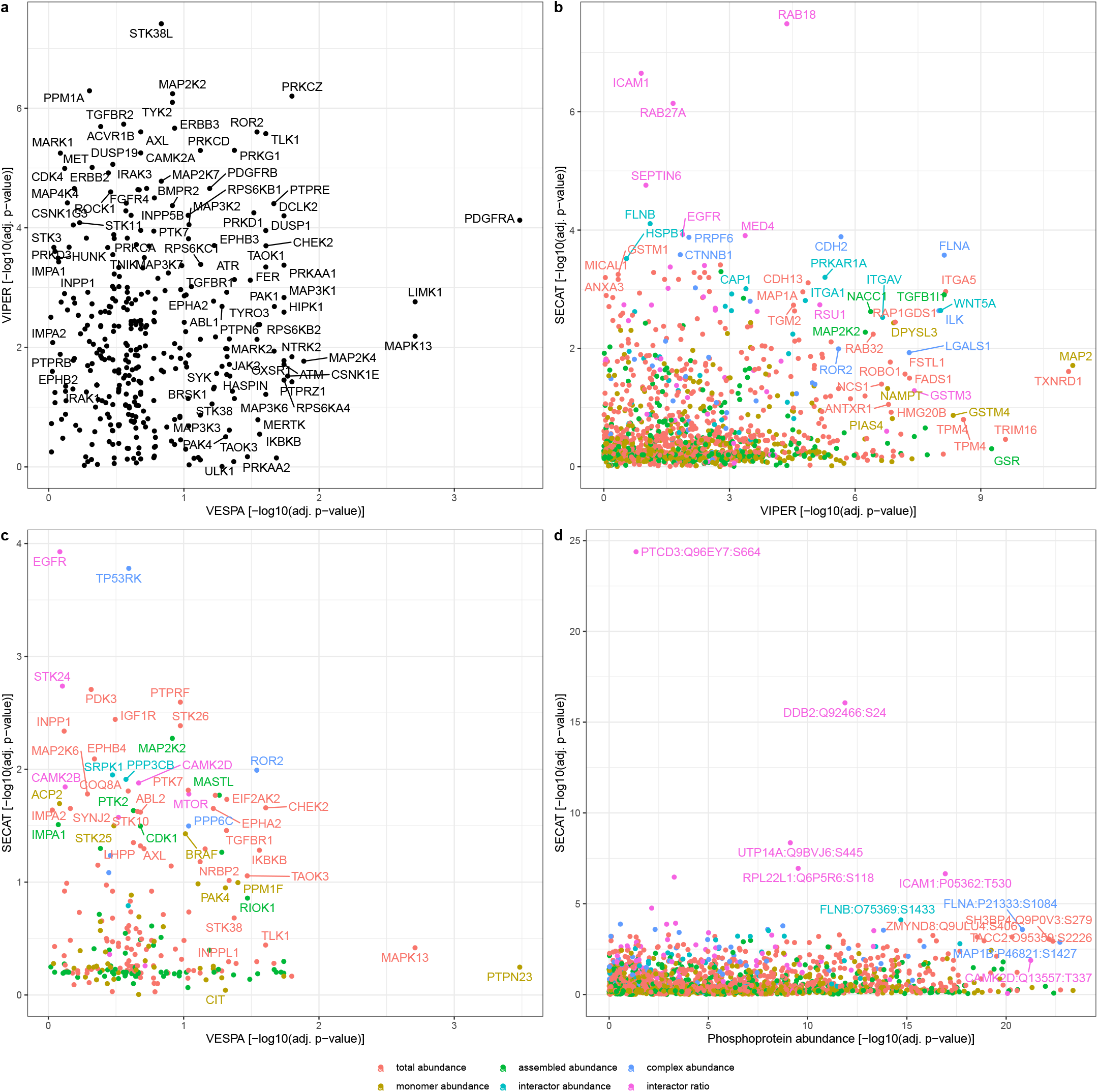
Correlation plot of differential VESPA, VIPER, SECAT protein activities and differential phosphopeptide abundances between HeLa Kyoto and CCL2 cell lines. **a)** Correlation plot between VESPA inferred signaling activity and VIPER inferred transcription factor activity. **b)** Correlation plot between VIPER inferred transcription factor activity and SECAT protein complex dynamics. **c)** Correlation plot between VESPA inferred signaling activity and SECAT protein complex dynamics. **d)** Correlation plot between phosphopeptide abundance and SECAT protein complex dynamics.

**Supplemental Figure 5.**
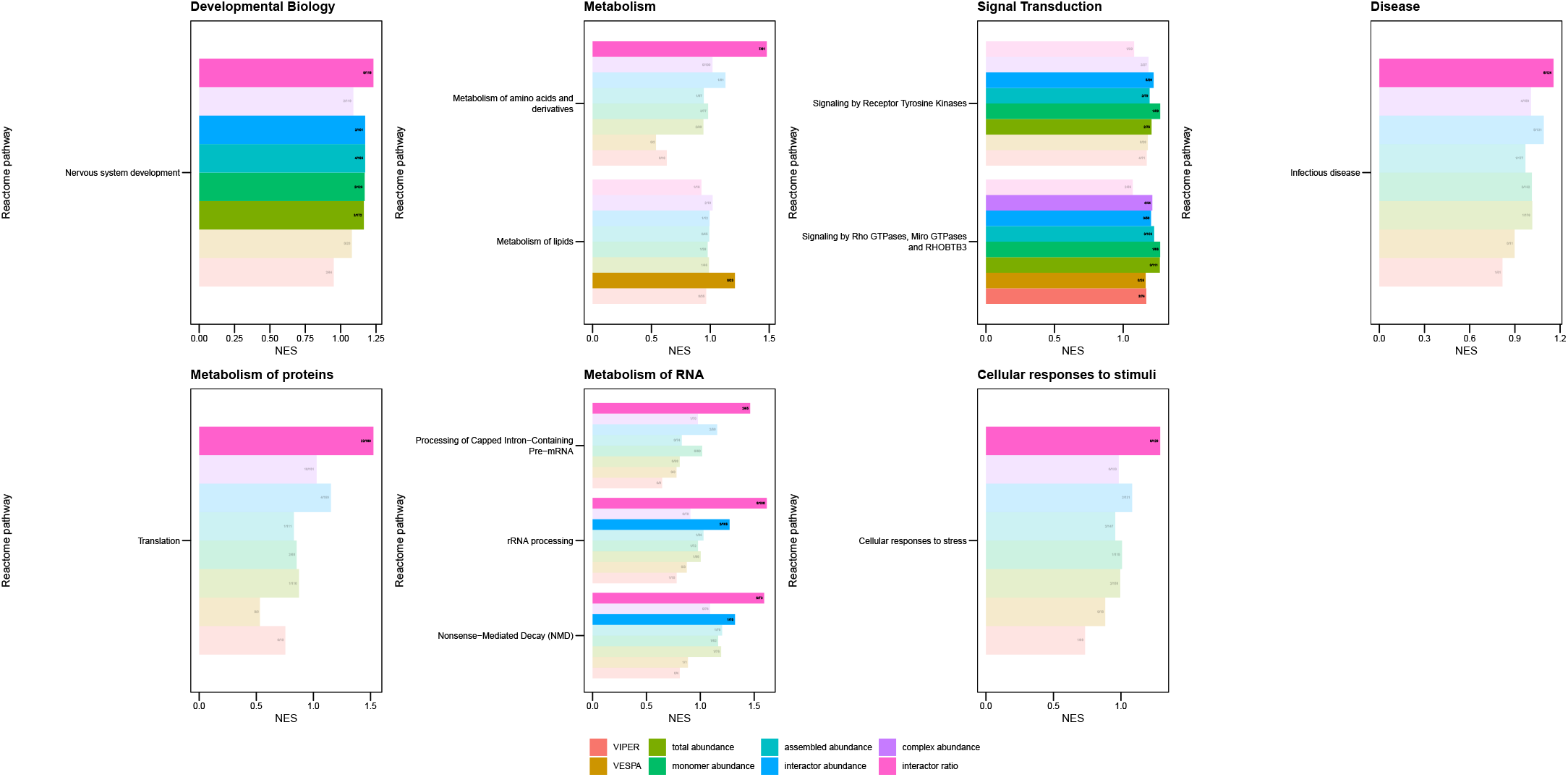
TieDIE network diffusion analysis with upstream VESPA layer. Opaque categories are significant (BH-adj. p-value < 0.05). The numbers in the barplots indicate [number of top 50 selected proteins part of the leading edge]/[number of proteins covered by upstream or downstream layer and present in leading edge].

**Supplemental Figure 6.**
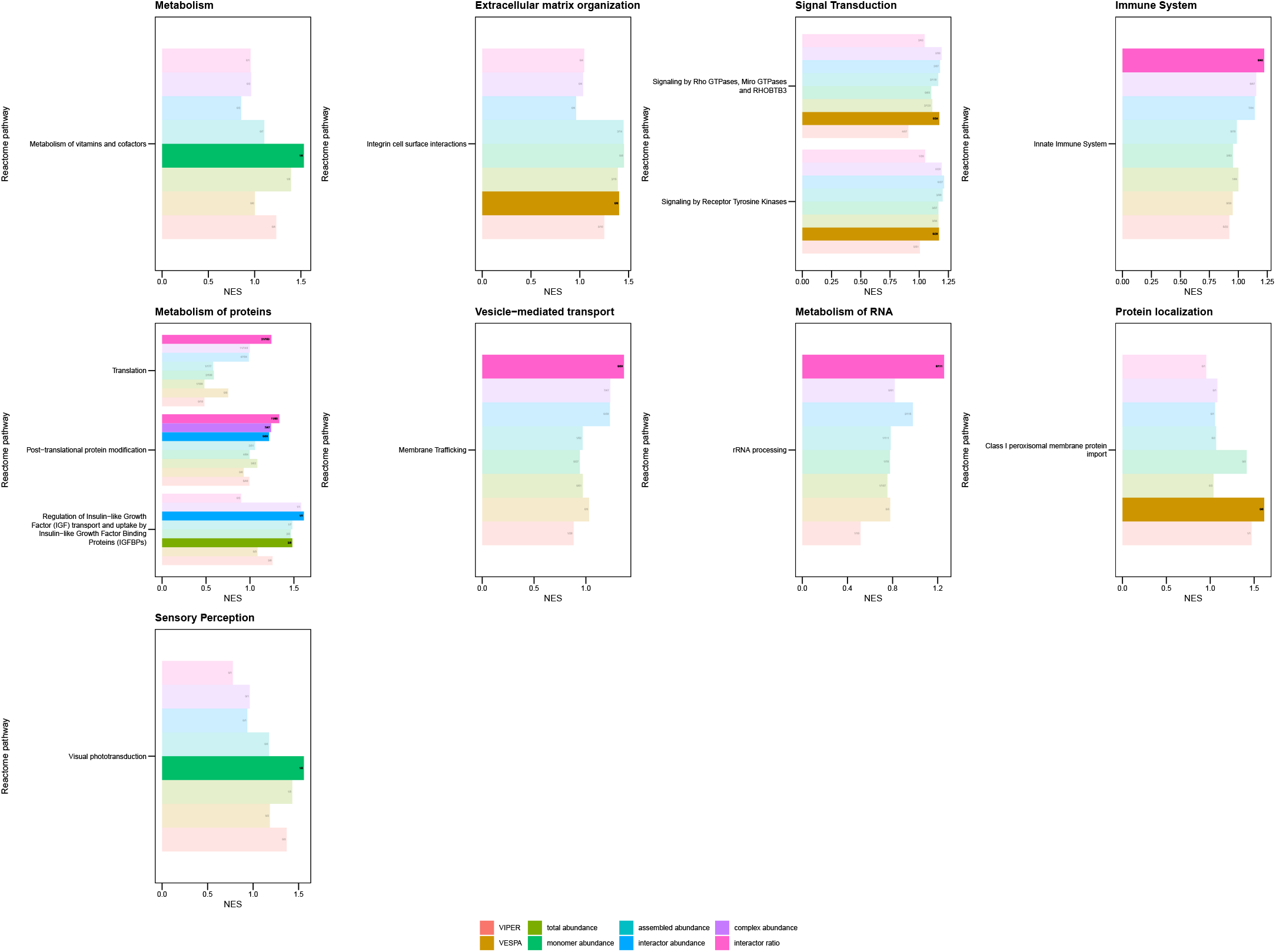
TieDIE network diffusion analysis with upstream VIPER layer. Opaque categories are significant (BH-adj. p-value < 0.05). The numbers in the barplots indicate [number of top 50 selected proteins part of the leading edge]/[number of proteins covered by upstream or downstream layer and present in leading edge].

**Supplemental Figure 7.**
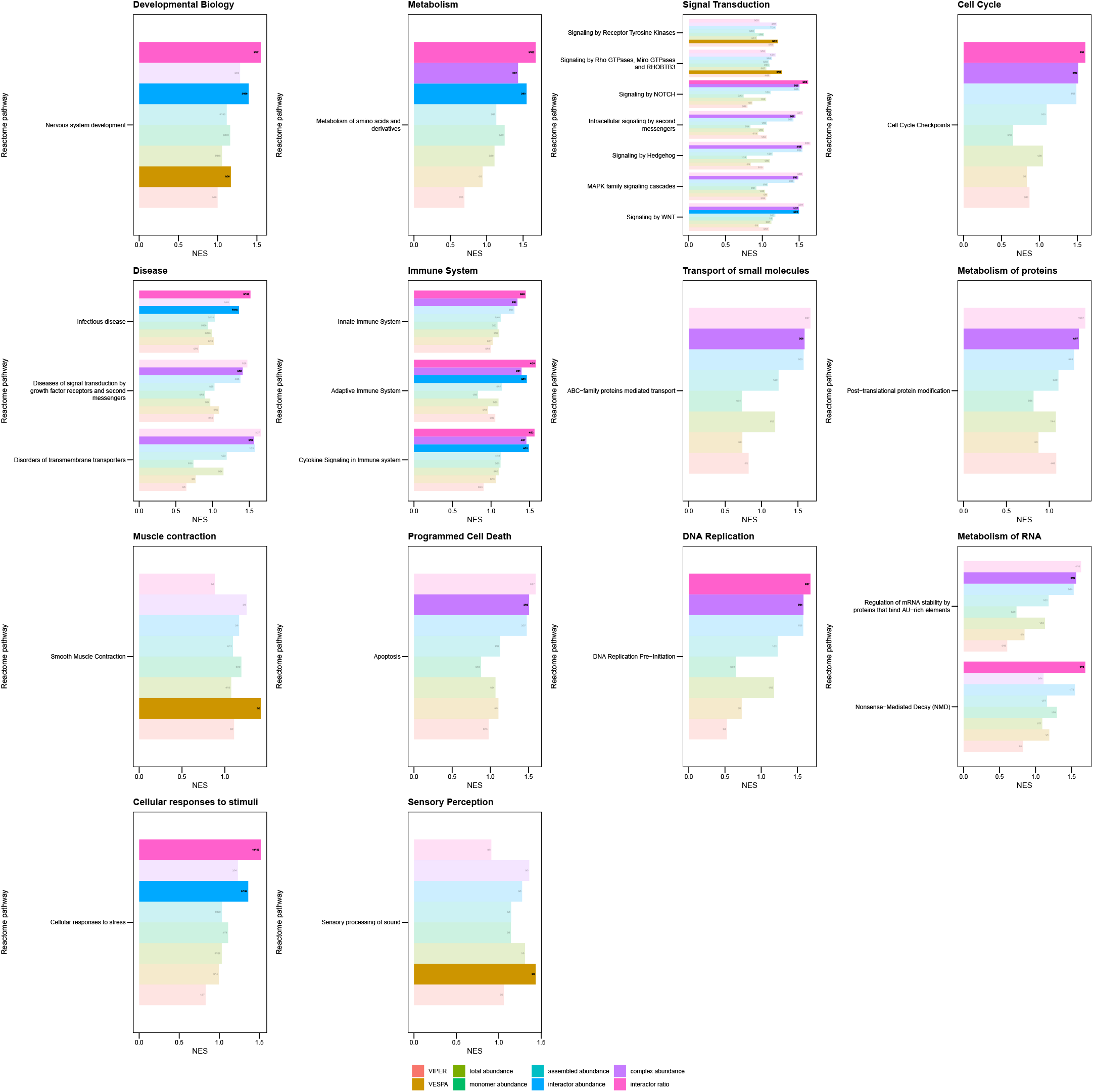
TieDIE network diffusion analysis with upstream SECAT total abundance layer. Opaque categories are significant (BH-adj. p-value < 0.05). The numbers in the barplots indicate [number of top 50 selected proteins part of the leading edge]/[number of proteins covered by upstream or downstream layer and present in leading edge].

**Supplemental Figure 8.**
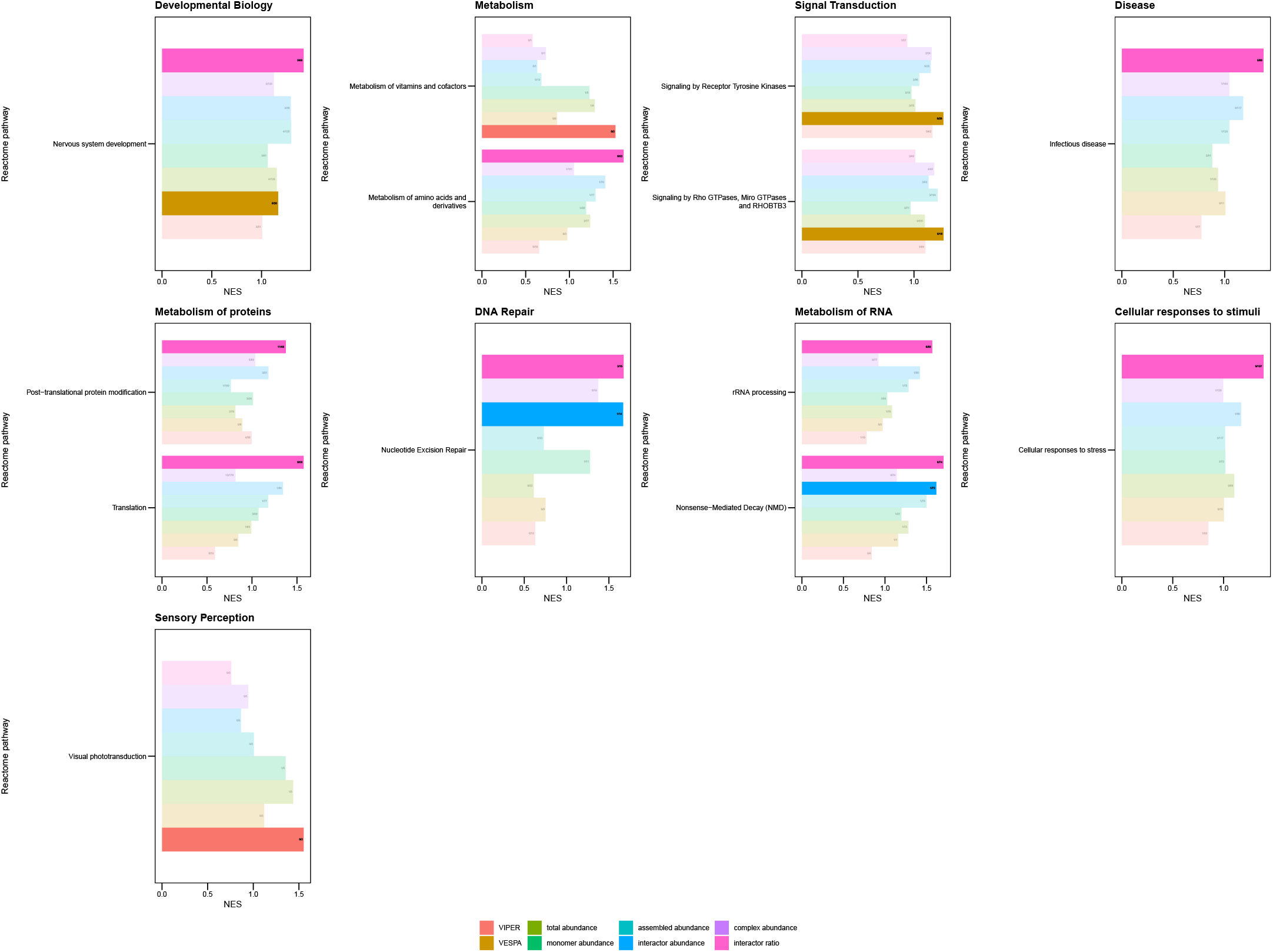
TieDIE network diffusion analysis with upstream SECAT monomer abundance layer. Opaque categories are significant (BH-adj. p-value < 0.05). The numbers in the barplots indicate [number of top 50 selected proteins part of the leading edge]/[number of proteins covered by upstream or downstream layer and present in leading edge].

**Supplemental Figure 9.**
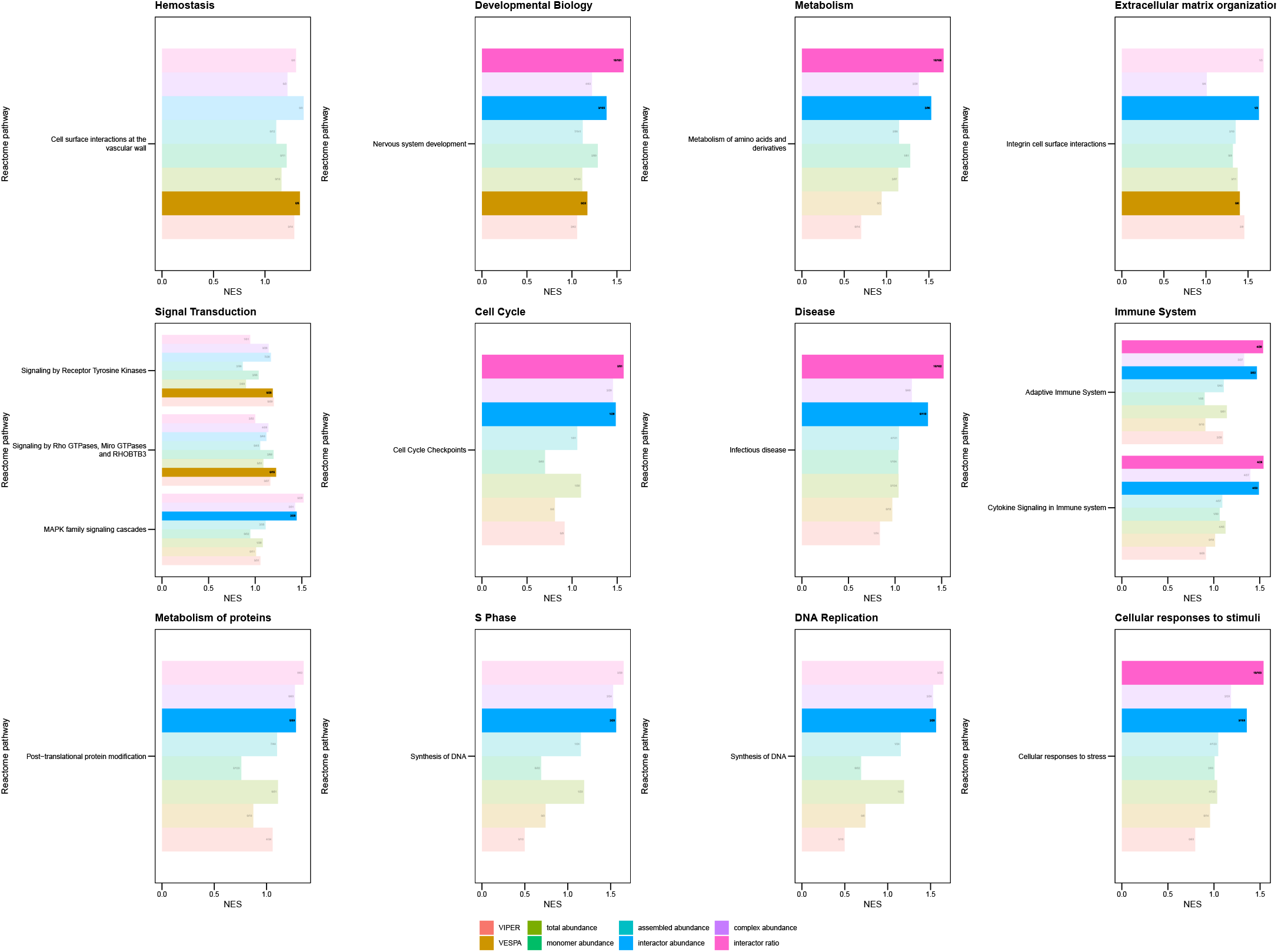
TieDIE network diffusion analysis with upstream SECAT assembled abundance layer. Opaque categories are significant (BH-adj. p-value < 0.05). The numbers in the barplots indicate [number of top 50 selected proteins part of the leading edge]/[number of proteins covered by upstream or downstream layer and present in leading edge].

**Supplemental Figure 10.**
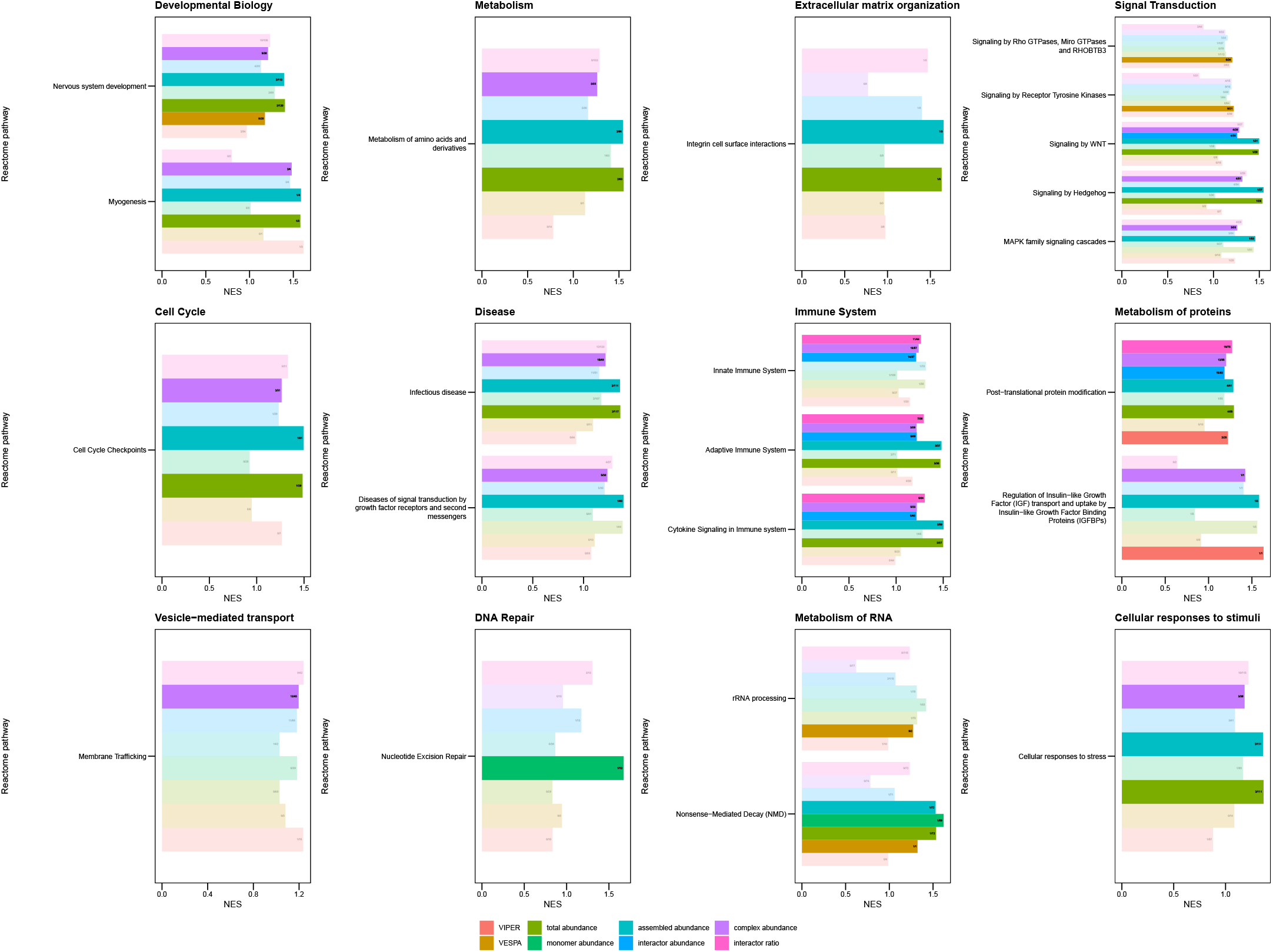
TieDIE network diffusion analysis with upstream SECAT interactor abundance layer. Opaque categories are significant (BH-adj. p-value < 0.05). The numbers in the barplots indicate [number of top 50 selected proteins part of the leading edge]/[number of proteins covered by upstream or downstream layer and present in leading edge].

**Supplemental Figure 11.**
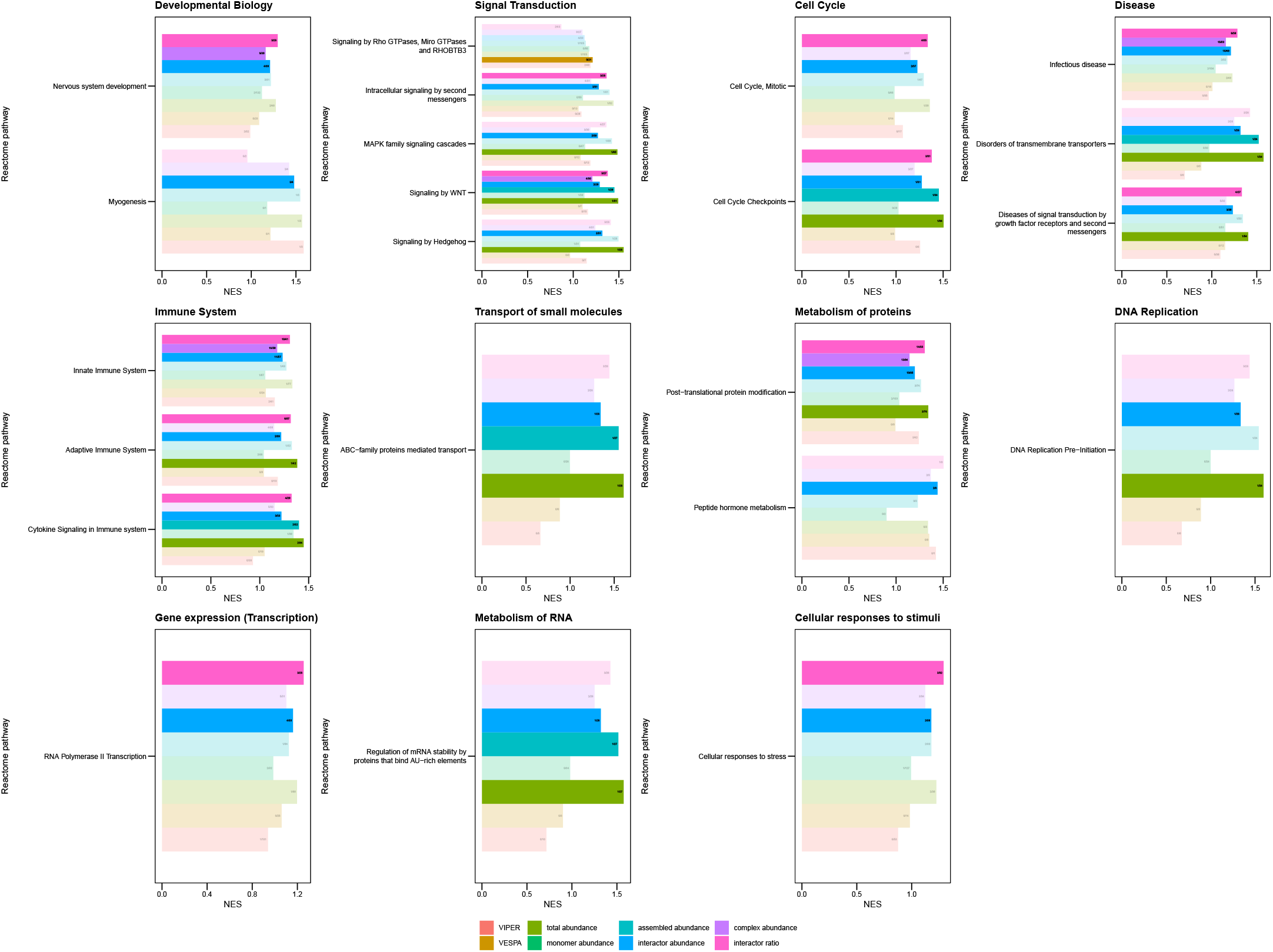
TieDIE network diffusion analysis with upstream SECAT complex abundance layer. Opaque categories are significant (BH-adj. p-value < 0.05). The numbers in the barplots indicate [number of top 50 selected proteins part of the leading edge]/[number of proteins covered by upstream or downstream layer and present in leading edge].

**Supplemental Figure 12.**
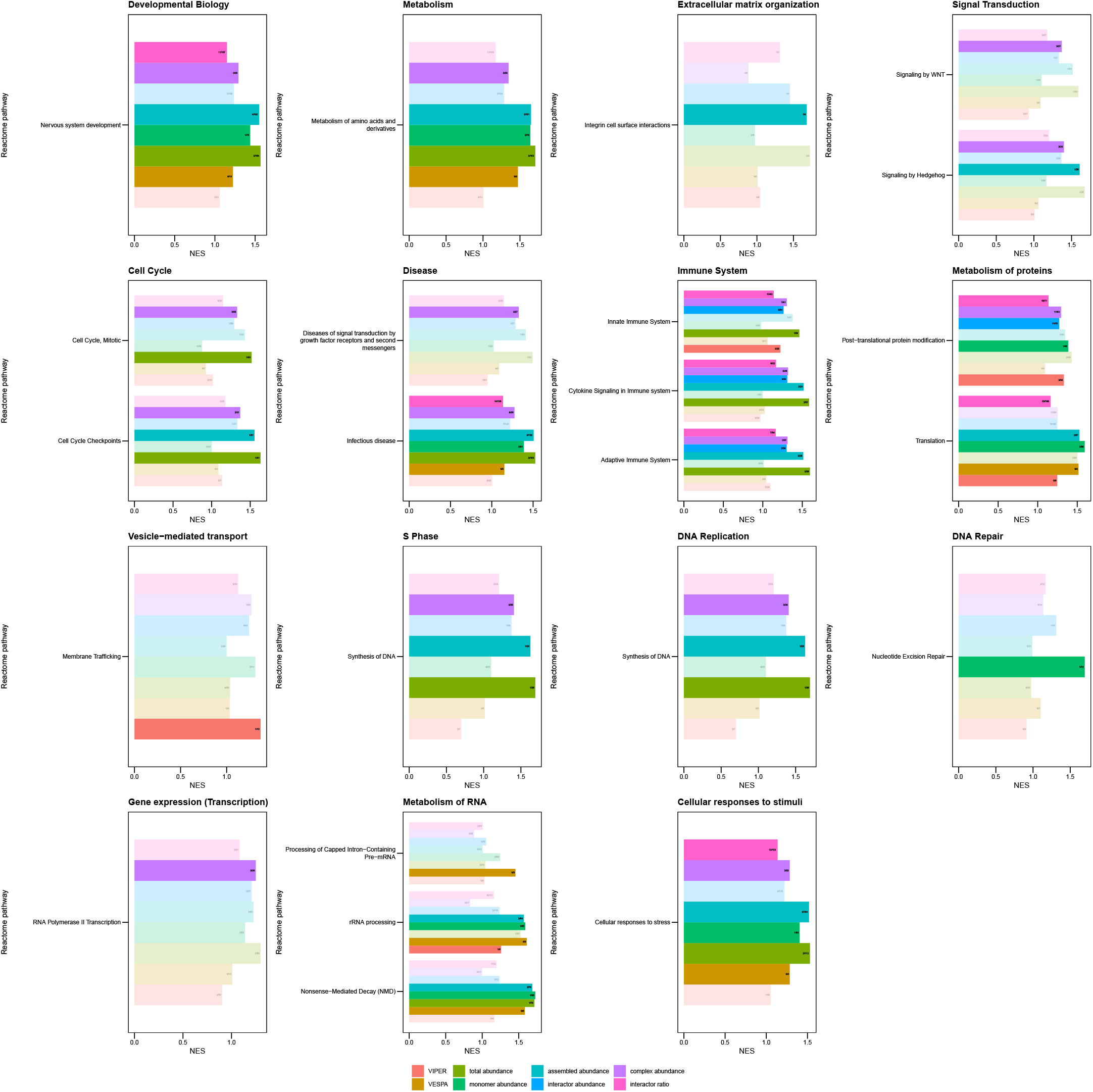
TieDIE network diffusion analysis with upstream SECAT interactor ratio layer. Opaque categories are significant (BH-adj. p-value < 0.05). The numbers in the barplots indicate [number of top 50 selected proteins part of the leading edge]/[number of proteins covered by upstream or downstream layer and present in leading edge].

**Supplemental Figure 13.**
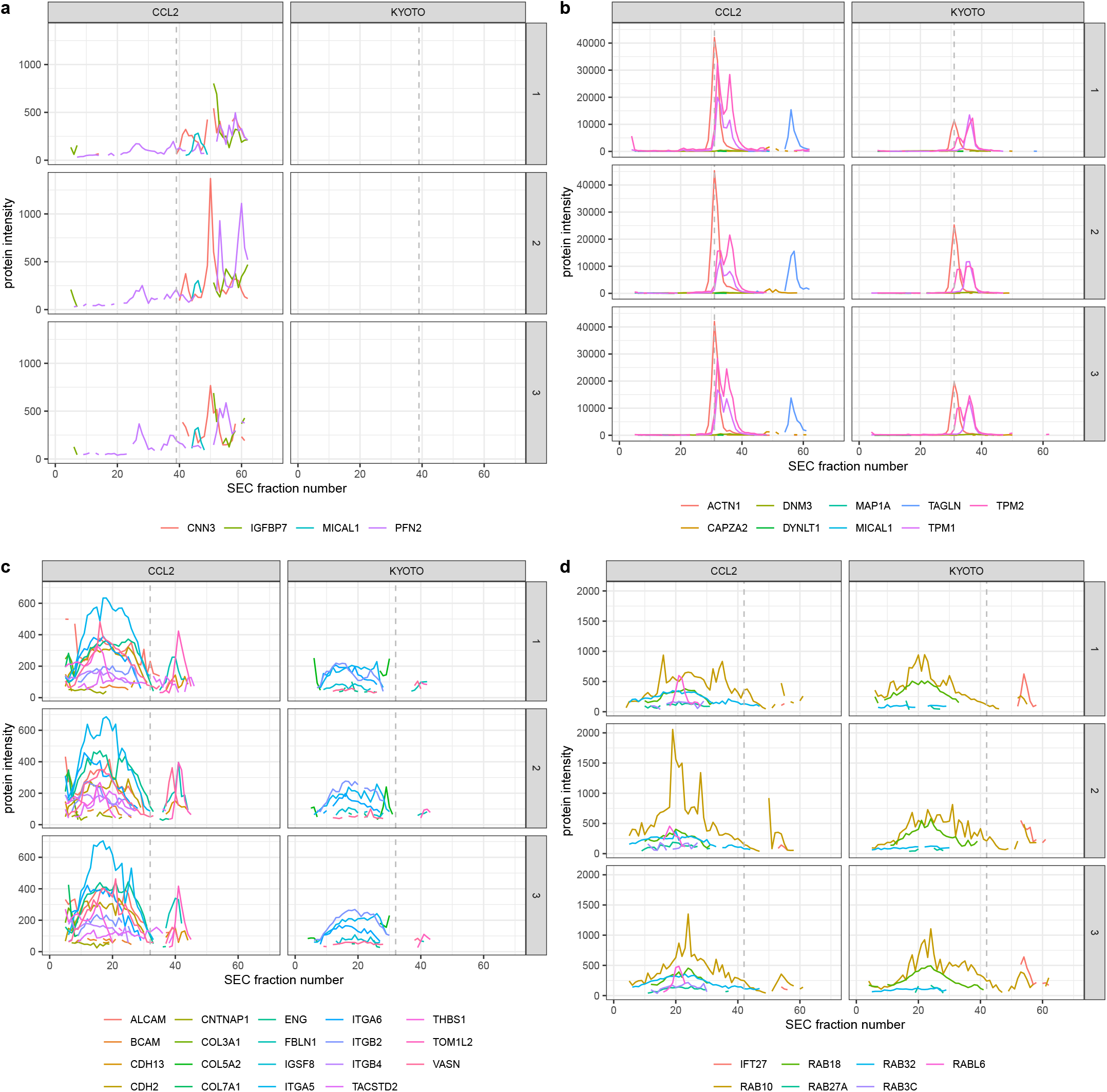
Supplemental protein-level SEC-SWATH profiles of HeLa CCL2 and Kyoto cell lines. Lines indicate different protein subunits, whereas the dashed line indicates the highest monomer threshold per group. **a-b)** Cytoskeleton-associated protein-level SEC-SWATH profiles of HeLa CCL2 and Kyoto cell lines. Lines indicate different protein subunits, whereas the dashed line indicates the highest monomer threshold per group. **c)** ECM- and cell surface-associated protein-level SEC-SWATH profiles of HeLa CCL2 and Kyoto cell lines. Lines indicate different protein subunits, whereas the dashed line indicates the highest monomer threshold per group. **d)** Rab-GTPase protein-level SEC-SWATH profiles of HeLa CCL2 and Kyoto cell lines.

**Supplemental Figure 14.**
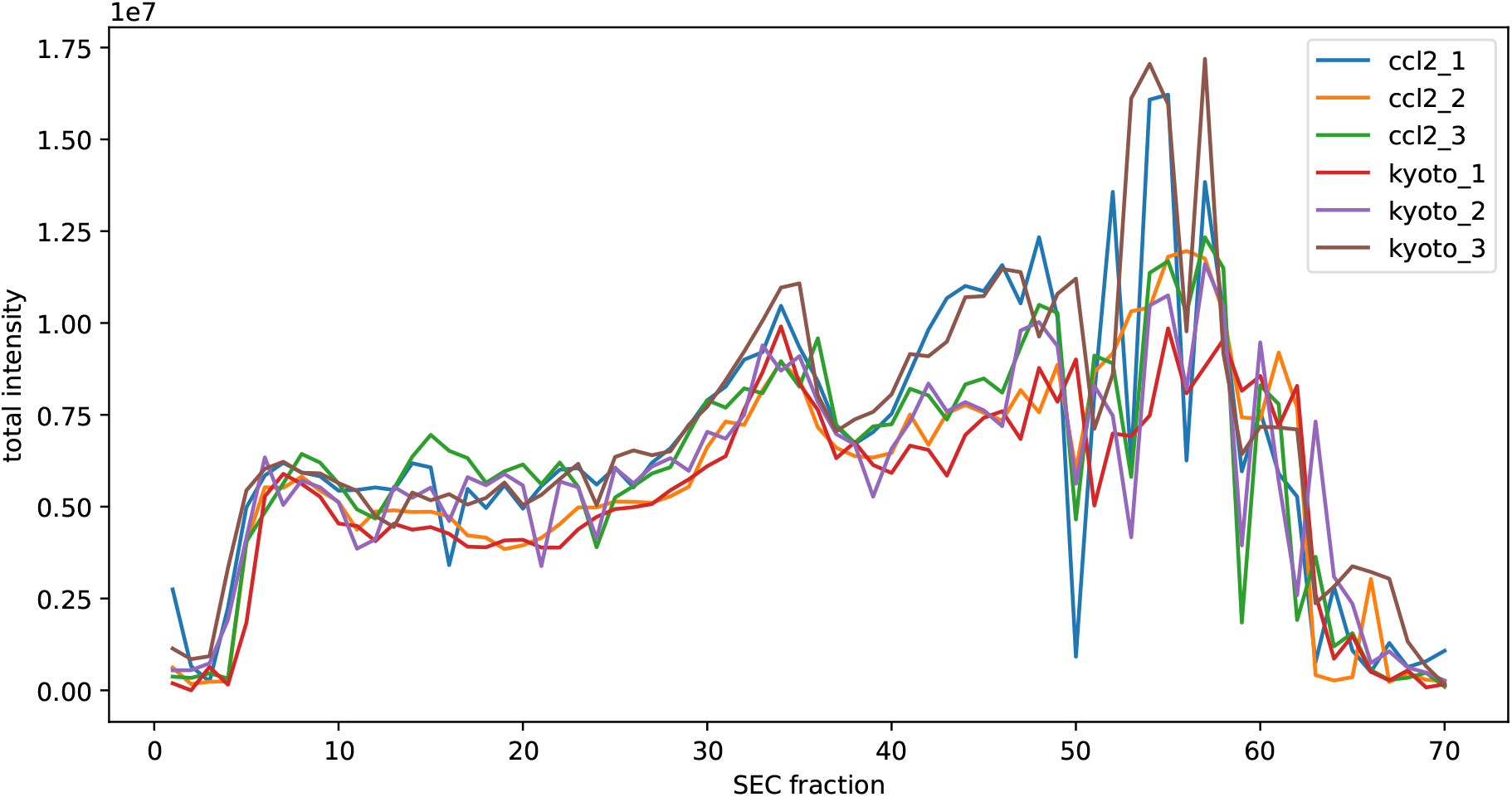
Summarized raw peptide precursor intensities over the SEC profile for the HeLa Kyoto vs. CCL2 data set.

**Supplemental Figure 15.**
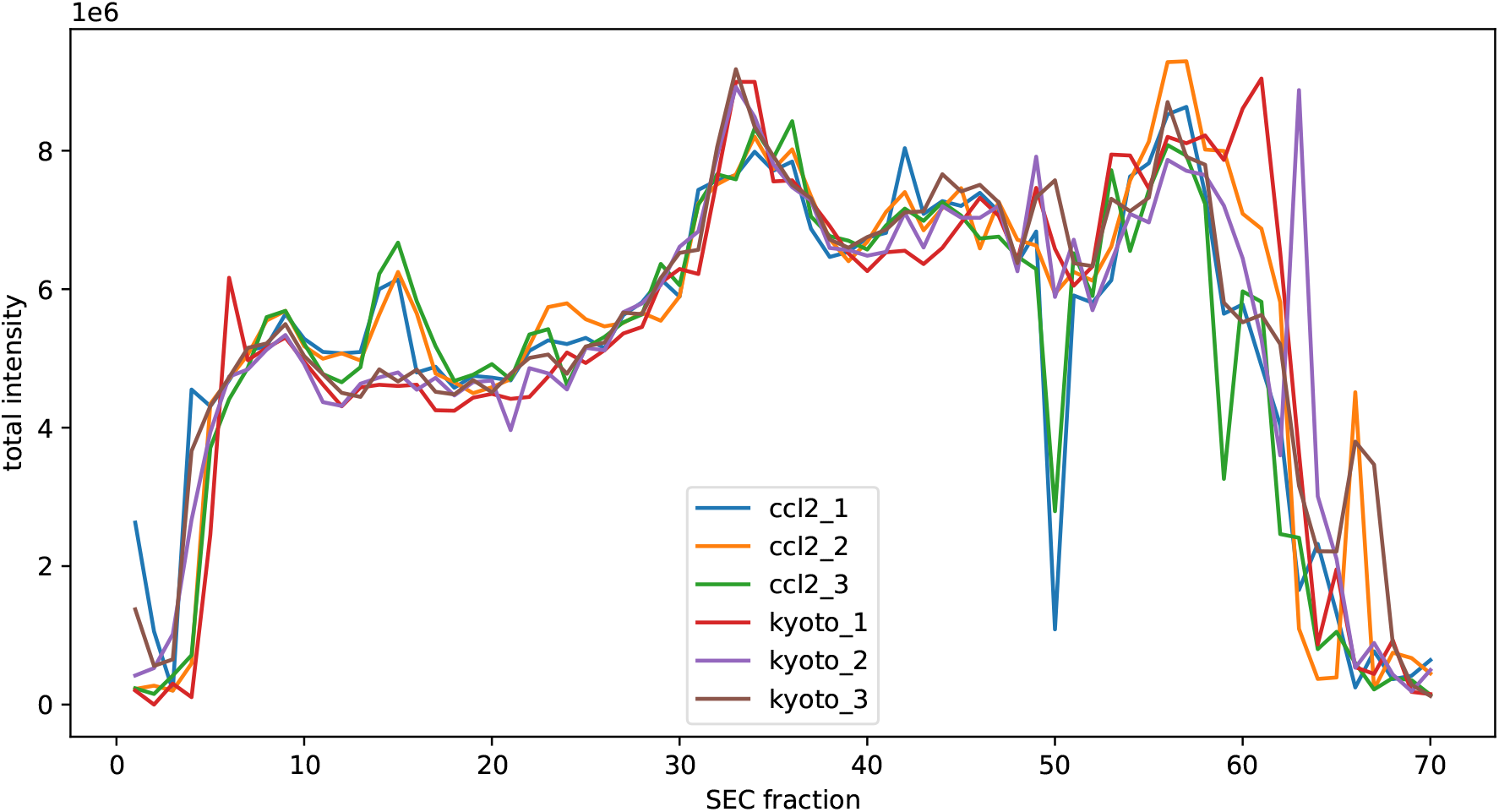
Summarized normalized peptide precursor intensities over the SEC profile for the HeLa Kyoto vs. CCL2 data set.

